# Cortical Reactivations Modulated by Local Inhibitory Circuits Mediate Memory Consolidation

**DOI:** 10.1101/2024.10.28.620563

**Authors:** Kristian Kinden Lensjø, Ingeborg Nymoen Nystuen, Frederik Sebastian Rogge, Kristin Tøndel, Arthur Sugden, Inga Shurnayte, Sverre Grødem, Anders Malthe-Sørenssen, Torkel Hafting, Mark L. Andermann, Marianne Fyhn

## Abstract

Highly salient events activate neurons across various brain regions. During subsequent rest or sleep, the activity patterns of these neurons often correlate with those observed during the preceding experience. Growing evidence suggests that these reactivations play a crucial role in memory consolidation, the process by which experiences are solidified in cortical networks for long-term storage.

Here, we demonstrate that reactivations in the lateral visual cortex are vital for the consolidation of visual association learning. By employing longitudinal two-photon Ca^2+^ imaging alongside paired LFP recordings in the hippocampus and cortex, we show that targeted manipulation of PV^+^ inhibitory neurons in the lateral visual cortex *after* daily training selectively attenuated cue-specific reactivations and learning, with no apparent effect on normal network function during training. In contrast, reactivations in the control group were biased towards salient cues, aligned with learning process and persisted for hours after training had ended. Overall, our results underscore a crucial role for cortical reactivations in memory consolidation.

## 1 Introduction

Memory consolidation enables the retention and integration of experiences, facilitating adaptive behaviors and survival. This consolidation process is proposed to rely on the concurrent activity of neurons in the hippocampus and cortex during offline periods, i.e., in resting periods and sleep following salient experiences [Squire et al., 2015, Buhry et al., 2011, Kaefer et al., 2022]. Within these offline periods, neurons that were active during the experience are reactivated in similar patterns [Wilson and McNaughton, 1994, Grosmark et al., 2021, Barry and Maguire, 2019, Foster, 2017]. Such reactivation events have been demonstrated in limbic areas (e.g., hippocampus and amygdala) and several areas of the cortex [Girardeau et al., 2017, Ji and Wilson, 2007, Chang et al., 2023] and are hypothesized to strengthen the connectivity between reactivated neurons. On a population level, sharp-wave ripples (SWRs) in the hippocampus and slower spindles in the cortex are often correlated in time during consolidation [Siapas and Wilson, 1998, Maingret et al., 2016]. This cortico-hippocampal coupling is believed to be important for the organization of activity across brain regions, supported by several observations that disrupting SWRs attenuates memory consolidation [Girardeau et al., 2009, Ego-Stengel and Wilson, 2009]. Importantly, cortical reactivations have been shown to occur during SWRs in both human and animal studies (e.g., [Sugden et al., 2020, Nguyen et al., 2024, Schreiner et al., 2024]). However, causal evidence for the importance of reactivations in consolidation largely remains elusive.

Emerging evidence indicates that the lateral visual cortex, and in particular the visual part of the postrhinal cortex (visPOR), plays an important role in learning visual associations. Here, the cortical activity changes across learning stages in different behavioral tasks, and inhibiting neuronal activity in visPOR significantly impairs task performance [Burgess et al., 2016, Ramesh et al., 2018, Goltstein et al., 2021]. Several lines of evidence from both human and animal studies show that the lateral visual cortex is involved in reactivations shortly after salient experiences [Deuker et al., 2013, Murty et al., 2017, Sugden et al., 2020]. Whether such reactivation events, immediately following salient experiences (e.g., Sugden et al. [2020]), reflect persistent activity involved in short-term memory or act as a component of consolidation is not clear. In comparison, ongoing activity in temporal association areas 24-72 hours after auditory-cued fear conditioning is essential for consolidating such memories [Grosso et al., 2015].

Recent work has suggested that inhibitory network control plays a role in memory consolidation [Ognjanovski et al., 2017, Mongillo et al., 2018]. The local network activity within the neocortex is under strong influence from inhibitory neurons that express parvalbumin (PV+) [Cardin et al., 2009, Sohal et al., 2009, Couey et al., 2013]. Here, the PV+ neurons reduce noise and spiking variability of excitatory output neurons [Atallah et al., 2012, Christensen et al., 2021, Nocon et al., 2023, Mongillo et al., 2018], suggesting an important role in coordinating activity at fast time-scales in moments of synchrony, such as reactivation events. Moreover, PV+ neurons have been implicated in shaping the local activity during sleep and in transitions between sleep states [Ognjanovski et al., 2017, Xia et al., 2017, Aime et al., 2022]. In particular, both Xia et al. (2017) and Ognjanovski et al. (2017) showed that reducing the activity of PV+ neurons after contextual fear conditioning prevented consolidation, where the manipulation of activity was accompanied by reduced SWR generation and cortico-hippocampal synchrony. However, more recent work from Abbas et al. [2018] showed that PV+ neurons did not influence long-range synchrony in a similar experiment but with more stringent manipulation of their activity. The contribution of PV+ neurons to cortico-hippocampal synchrony during memory processing is still unclear.

Given the precise reactivation patterns of cell ensembles previously observed after learning, their tight coupling to hippocampal SWRs, and the importance of PV+ neurons in local cortical circuits, we hypothesize firstly that cortical reactivations are essential for memory consolidation and, secondly, that PV+ inhibitory neurons provide the tight network regulation and temporal control required for accurate reactivation patterns. To test this, we trained mice in a visual Go/NoGo task where learning occurs in a gradual manner across many days [Sugden et al., 2020]. We tracked neuronal activity using two-photon Ca2+ imaging in the lateral visual cortex, centered on visPOR, before, during, and after daily training. To directly assess the synchrony between the hippocampus and cortex, we conducted paired recordings of local field potential in visPOR and CA1 of the hippocampus in post-training rest. Building on previous studies that have silenced or lesioned cortex during memory consolidation (e.g., Sacco and Sacchetti [2010], Grosso et al. [2015]), we aimed to specifically target reactivations by transient manipulation of neural activity. Importantly, our approach was designed to achieve this without reducing the activity of excitatory neurons in the cortex. To this end, we reduced the activity of PV+ neurons in visPOR in every other session (day) in the offline state, i.e., during quiet waking after completion of daily training. Remarkably, we found that partial inhibition of PV+ cells impaired across-day learning. The behavioral performance improved following sessions in which the network was intact, only to revert to chance levels when the subsequent session involved reducing PV+ activity. This pattern indicates that the manipulation induced amnesia for the learning acquired on the previous day. Within the lateral visual cortex, we observed that the learning-induced changes seen in controls - such as the development of reward-cue bias in cortical responses - were completely prevented by the partial offline inhibition of PV+ neurons. Importantly, the activity manipulation did not affect the response selectivity or overall fraction of cue-responsive cells that is inherent to the lateral visual cortex [Burgess et al., 2016]. Finally, in the offline activity of controls during quiet waking, we observed a transient bias in the incidence rates of reactivations towards the most salient, rewarded cue. In contrast, we show that offline inhibition of PV+ neurons leads to an overall increase in spontaneous activity but a strong reduction in both the incidence and specificity of reactivations to salient cues. Altogether, our findings demonstrate that reactivations in the lateral visual cortex, which occur immediately following salient experiences, depend on intact inhibitory activity to mediate memory consolidation.

## 2 Results

### Offline reduction of PV+ activity prevents learning

To investigate how the structure of offline activity affects memory consolidation, we expressed the inhibitory DREADD receptor hM4Di in PV+ neurons using a viral vector (AAV5-hSyn-DIO-hM4Di-mCherry, AAV-PHP.eB-hSyn.Flex-mRuby3 in controls) in PV-Cre mice. Bilateral virus injections (150 nL per hemisphere) were targeted to the lateral visual cortex, specifically visPOR (Fig. 1A, see Supplementary Fig. 1a for extent of the spread of injections). Food-restricted (85 − 90%) animals were then trained in an operant Go-NoGo visual discrimination task [Burgess et al., 2016, Ramesh et al., 2018, Sugden et al., 2020] to learn three separate cue-outcome associations in order to acquire food reward (plus cue, 0°, Ensure) and avoid punishment (minus cue, 270°, 0.01M quinine). A third cue with no outcome was also included to have a neutral condition (135°) with no saliency (Fig. 1B). At the end of daily training sessions, i.e. in the offline state, the mice received a single 125 *µ*L intraperitoneal injection of either 0.9% saline solution or Clozapine N-oxide (CNO, 5 mg/kg body weight, diluted in 0.9% saline) (Fig. 1B, right panel), where CNO was injected every other day (even and odd sessions, counterbalanced between individuals).

**Figure 1:**
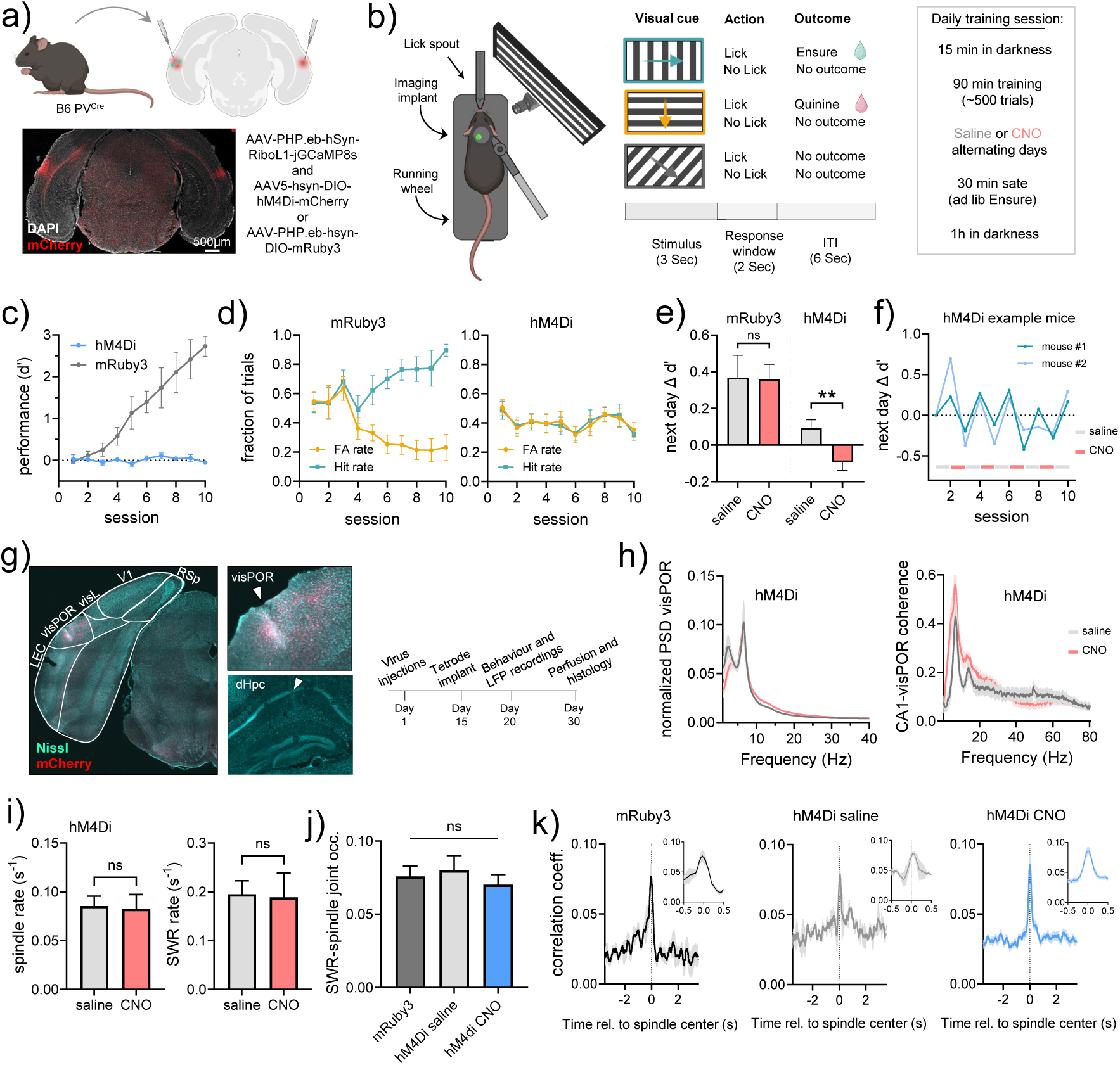
Transient inhibition of PV+ cells after training prevents consolidation. **a)** Schematic of virus injection sites in PV-Cre mice. b) Overview of experiment and daily training protocol. Note that CNO or saline was only injected *after* training had been completed for the day. c) Behavioral performance (d’) for mice injected with mRuby3 (n=7) or hM4Di (n=13) in visPOR across 10 days of training. The difference was statistically significant p<0.0001, generalized linear mixed-effects model (GLMM). d) False alarm and hit rates across training. e) Relative change in task performance (d’) on a given session following either saline or CNO injection in the previous session (day). The performance of hM4Di mice decreased the day after CNO injection (mRuby3 p=0.96, hM4Di p=0.0052 (two-tailed unpaired t-test). f) Relative change in performance (d’) from day to day for two example mice (hM4Di) receiving CNO or saline on alternating days (red and grey bars). g) Location of LFP recording electrodes in CA1 and visPOR, and experimental timeline. Left: coronal section showing the spread of the hM4Di expression in one hemisphere, restricted to visPOR. Middle: tetrode tracks in visPOR (upper) and CA1 (lower). h) Mean power spectral density (PSD) in visPOR and coherence of LFP signals between CA1 and visPOR. No significant differences between sessions were observed (only shown for internal comparison within hM4Di-injected mice). i) Incidence rates of cortical spindles and hippocampal SWRs. j) Joint occurrence rate of isolated SWRs and spindle events from CA1 and visPOR, respectively. No significant difference between groups was found. One-way ANOVA, F=0.32, p= 0.72. k) Cross-correlation between spindles and SWRs, shown ±3 and 0.5 (inset) seconds from spindle center. All graphs represent mean± s.e.m.

Control mice with expression of inert mRuby3 (n=7) were able to learn the task to a high-performance level (>90 % correct, shown as discriminability, d’) within 10 days (Fig. 1C-D, the individual behavioral performance shown in Supplementary Fig. 1b). Strikingly, animals injected with hM4Di (n=13) did not show any sign of improved performance across the training period, despite the fact that the performance often improved during training sessions (Supplementary Fig. 1c). When we compared the performance from day to day in mice injected with saline or CNO after the previous training session, we observed no effect of CNO compared to saline injections in mRuby3 control mice (Fig. 1E). In contrast, in hM4Di-injected animals, performance improved in the training sessions where saline was injected following the prior session, and it decreased by a similar amount in training if CNO was injected the previous day. These opposite effects of saline and CNO sessions could be observed in individual animals, and the performance consequently remained close to chance levels (Fig. 1F). This suggests that the hMDi activation induced amnesia for the information acquired during the session, preventing the consolidation of learning that would have otherwise occurred following the saline injection.

To test whether offline PV+ inactivation simply delayed learning, a subset of animals were trained for an extended period with CNO injections every second day (up to 25 days). The performance of these mice showed no improvement, despite undergoing more than twice the number of training sessions (Supplementary Fig. 1d). Moreover, animals with hM4Di injected into only one hemisphere displayed behavior comparable to that of the mRuby3-injected control group (Supplementary Fig. 1e. Importantly, the behavioral effects of offline hM4Di activation experiments were reproduced in two labs (Supplementary Fig. 1f). The animals’ behavior during training, including overall lick rate and pre-training body weight, did not differ from that of the mRuby3-injected controls (Fig. 1D, Supplementary Fig. 1g).

Consolidation of memories depends on communication between different brain areas. To test how the inactivation of PV+ neurons in visPOR affected population activity and synchrony with other regions of the brain, we performed extracellular recordings of the local field potential (LFP) from the dorsal hippocampal CA1 and the visPOR in the same hemisphere (Fig. 1G). Tetrodes were implanted, and simultaneous recordings of the LFP from both brain areas were performed after daily training while mice rested on the running wheel in darkness. Virus injections and saline and CNO injections were performed as described above. We first computed the power spectral density and coherence across the two brain areas at the lower end of the frequency spectrum. No significant effects of hM4Di activation were observed (Fig. 1H). Next, we identified time periods in which SWRs or cortical spindles occurred, using established criteria (e.g., Xia et al. [2017]). We found no effect of hM4Di activation on the incidence rates of hippocampal SWRs or spindles in visPOR, compared to recordings from the same animals with saline injections (Fig. 1H, Supplementary Fig. 1h). However, when compared to mRuby3-injected controls, we observed a small increase in the incidence rate for cortical spindles (Supplementary Fig. 1h). To examine whether manipulating the PV+ neuron population in visPOR influenced the synchrony between SWRs and cortical spindles, we first calculated the joint occurrence rate, defined as the frequency at which SWRs occurred within a ±25 ms window from each spindle center [Maingret et al., 2016]. In contrast to previous work employing similar methodology in other brain regions, such as CA1 and the anterior cingulate cortex [Xia et al., 2017], we found no differences between the groups (Fig. 1J). To investigate this in greater detail, we computed the cross-correlation between SWRs and cortical spindles occurring within the same time frame (± 4 sec from the spindle center). Our results demonstrated that, although somewhat weak, the synchrony between the two areas was comparable to earlier work from other brain areas (e.g. Siapas and Wilson [1998], Maingret et al. [2016]), with a significant increase in synchrony around the spindle center. This held true across all groups (hM4Di saline and CNO sessions, and mRuby3) and suggested that the cortico-hippocampal synchrony was not affected by the local manipulation of the visPOR (Fig. 1K).

A potential confounding factor in experiments involving reduced activity of inhibitory neurons is the increased risk of epileptiform activity. To this end, we performed a similar analysis as those described in [Steinmetz et al., 2017]. We detected no epileptiform event or seizure-like activity in any of the recordings (Supplementary Fig. 2), which is in line with previous work on moderate suppression of PV+ activity [Atallah et al., 2012].

### Learning-induces changes in cortical neurons are prevented by offline reduction of PV**+** activity

After confirming that offline inhibition of PV+ neurons in visPOR prevented learning with no apparent effect on cortico-hippocampal synchrony, we next investigated how the cortical network responded during different training phases. To achieve this, we conducted longitudinal two-photon imaging to record the activity of large populations of neurons across the training period. In order to maximize the number of neurons sampled and reduce contamination from the neuropil activity, we expressed the Ca2+ indicator RiboL1-jGCaMP8s [Grødem et al., 2023] in all cells using the general neuron promoter hSyn, together with hM4Di or mRuby3 targeted to PV+ neurons as described above (Fig. 1A and Fig. 2A-B). Expression of the viral constructs was confirmed with *in vivo* widefield imaging and *post-mortem* histology (Fig. 2A, Supplementary Fig. 3a). We recorded from neurons in layer 2/3 (120-250µm below the cortical surface) from eight mice (n=4 injected with mRuby3 and n=4 injected with hM4Di) starting from the first naive training session and throughout the training period to expert performance (total number of identified cells: 42977, mean number of cells per mouse per session: 588±302 (mean±standard deviation), Supplementary Fig. 3c, and Supplementary Fig. 4a). Data from these eight mice were used in all subsequent analyses of training and offline activity. Each recording session consisted of a brief baseline recording in darkness, followed by 500 training trials across three 30-minute runs and 1 hour post-training rest in darkness (Fig. 1B). Saline or CNO was injected directly after training. This was followed by a 30-minute period with *ad libitum* access to Ensure to promote quiet waking and thus increase the likelihood of detecting reactivation events [Sugden et al., 2020], before we performed the post-training rest recordings. We first focused our analysis on the daily training runs when mice were presented with visual cues. Neurons were classified as visually responsive if they exhibited a significant increase in relative fluorescence during the first two seconds of the stimulus window to any cue, as compared to a one-second baseline preceding cue presentation. Cue bias was assigned based on which cue elicited the strongest response. This simple classification revealed three distinct ensembles with different orientation preferences (example of pairwise correlations across the sorted cue-responsive population shown in Supplementary Fig. 4c). Earlier reports have indicated that approximately 30% of neurons in visPOR are visually responsive with some orientation selectivity in naive mice [Burgess et al., 2016], and that neurons in this area respond to both stimulus identity, rewards and the combination of stimulus and reward [Ramesh et al., 2018, McGuire et al., 2022]. We observed a comparable fraction of visually responsive neurons in our data for both groups of animals in the very first training session (Supplementary Fig. 4b and d). When we compared the responses to the visual cues over the course of the experiment, we observed a large shift towards the “plus” cue presentations in mRuby3-injected controls, while no such development was observed in hM4Di-injected mice (Fig. 2B, Supplementary Fig. 5a). To investigate this in greater detail, we defined three phases of the training period: naive (first and second session), training (sessions with the highest shift in behavioral performance, sessions 5 and 6 in two animals, 7 and 8 in two animals) and late (final two sessions). For the hM4Di injected mice, where no change in performance was observed, we used sessions 5 and 6 as the “learning” phase data. When we compared the responses to the three cues across different stages of training in mRuby3-injected controls, we found a significant increase in the fraction of visually responsive cells with a bias towards the “plus” cue compared to both the “minus” and neutral cues (Fig. 2C, left panel). The large increase in cue bias appeared in the learning stage and persisted through the late stages of training. This was also true when comparing the development across learning stages within the entire recorded population and not limited to cue-responsive cells (Supplementary Fig. 4d). When focusing our analysis solely on the stimulus responses occurring within the first second of cue presentation and prior to any licking behavior, we continued to observe the same trend in relation to learning (Supplementary Fig. 4e). However, this observation did not reach statistical significance. This suggests that the heightened responses observed in the mRuby3-injected controls were a result of both an enhanced response to the plus cue itself and the successful combination of cue and reward pairing. In contrast, in hM4Di-injected mice, no development in the cue bias was detected across training stages (Fig. 2C, right panel). Notably, the proportion of neurons showing significant response to any visual cue remained stable in both groups (Supplementary Fig. 3b), indicating that the cortical network stayed intact during training sessions in hM4Di-injected mice. However, this fraction did not change across training in the cue-outcome association task, likely due to the manipulation of PV+ cells during the post-training period.

**Figure 2:**
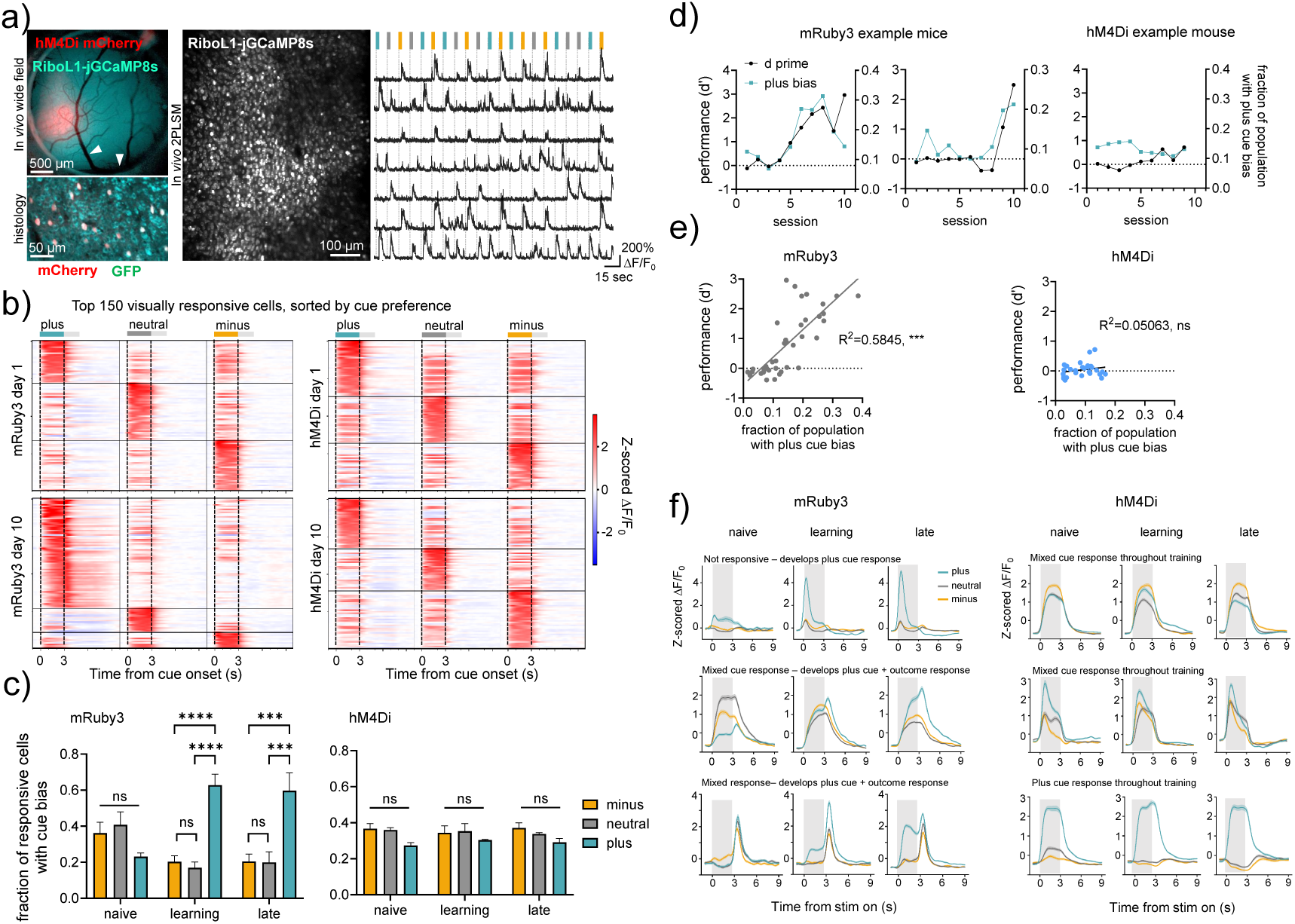
Cortical network changes with learning are prevented by offline hM4Di activation. **a)** Representative example from one imaging mouse expressing hM4Di in PV^+^ neurons and RiboL1-jGCaMP8s in all neurons. Expression of DIO-hM4Di-mCherry and the Ca^2+^ indicator RiboL1-jGCaMP8s was confirmed by *in vivo* widefield imaging and post-mortem histology. Right panel: changes in fluorescence during visual stimulation, selection of cells from the two-photon microscopy example field-of-view. Color bars indicate stimulus identity. **b)** Color-coded PSTH from the top 150 visually responsive cells on the first and last day of training from one representative mouse per group. In both groups, the cue preference was evenly distributed in naive mice. While mRuby3 animals developed a strong bias for the rewarded “plus” cue, a similar change was not detected in hM4Di-injected animals. **c)** Quantification of cue-bias across training phases normalized to the fraction of the population responding to any cue, p<0.0001 (minus vs. plus, “learning”), p<0.0001 (neutral vs. plus, “learning”), p= 0.0001 (minus vs. plus “late”), p= 0.0001 (neutral vs. plus, “late”), generalized linear mixed-effects model with post-hoc Tukey’s multiple comparisons tests. For the hM4Di mice, no significant differences were observed. **d)** Behavioral performance (d’, grey line) and the fraction of cells with a “plus” cue bias (green line) from representative mice in both groups. **e)** Quantification of the relationship between behavioral performance and “plus” cue bias. Linear regression, R^2^=0.5845, p<0.0001, n=4 mice (mRuby3). In hM4Di mice, we found no significant relationship. **f)** Examples of unique single cells tracked across the experiment, shown as the mean response to the first 10 cues on a given session from stimulus onset (*t* = 0). Line color indicates stimulus identity.

If the behavioral changes were, at least in part, a result of the development of cortical cue bias, we would anticipate that these changes would correlate with the individual learning curves of the animals. Indeed, in mRuby3-injected controls, we found a strong relationship between the fraction of cells exhibiting a “plus” cue bias and task performance (Fig. 2D and E). In hM4Di-injected animals, however, there was no development in either plus cue bias or performance.

In addition to examining the population-level responses during training, we investigated single-cell dynamics. This analysis was critical as learning-induced changes at the individual cell level might be masked by the aggregate population activity. We focused on single cells that were consistently tracked throughout the entire experimental period (Supplementary Fig. 5b). In mRuby3-injected controls, we observed a wide range of dynamic response profiles, where many cells gained response bias to the plus cue or reward (examples shown in Fig. 2F, left panel). In hM4Di-injected mice, we observed visual responses and cue selectivity that remained stable across the experiment but with no apparent changes in single-cell responses related to training. These findings support the notion that the cortical network remained in a “naive”-like state despite training, as the offline consolidation was prevented by the hM4Di activation.

### Cortical reactivations promote memory consolidation

Given that we observed normal visual responses but no apparent learning with behavioral training, we next focused on the offline population activity during post-training rest (i.e., after injections of either saline or CNO and a period of *ad libitum* consumption of Ensure) (Fig. 1B). Cue-specific cortical reactivations of cellular activity patterns in a time-compressed manner during resting states are hypothesized to play a crucial role in memory consolidation (e.g. Sugden et al. [2020]). This process relies on tight coupling within and between brain areas and requires precise regulation by the inhibitory network.

To detect cue-specific reactivations in visPOR, we trained and tested the performance of three different classifiers (Logistical regression, QDA, and Random Forest) using data from training sessions. For every session from each mouse, we trained the classifiers on a random selection of two thirds of the baseline activity in darkness (before training) and during training. The classifiers made predictions based on the recorded activity in the remaining one third of the data set, which were compared to the timing and true stimulus identity. The method with the overall best performance was the Random Forest classifier, which classified the stimulus and non-stimulus events with high accuracy (mean prediction accuracy of 98, 7%) and a low rate of false positives (<1%) (Supplementary Fig. 6d). When shuffling the data in time (on a frame-by-frame basis) and by cell ID prior to predicting visual responses in the test set, the prediction accuracy dropped to less than 5% (Supplementary Fig. 6e). This classifier was subsequently applied to all training data sets from each mouse to predict outcomes based on the recordings obtained during the post-training rest period. Periods where the animals were moving were excluded. One effect of reducing the activity of inhibitory neurons could be that the spontaneous activity of nearby excitatory neurons increases, thereby increasing ‘noise’ in the data. To maintain a conservative definition of reactivations, we restricted our analysis to periods when synchronous events occurred within the population of cue-responsive cells (similar to Nguyen et al. [2024]). The events identified by the classifier as stimulus-related activity patterns during these periods were defined as reactivations. As shown in the example events in Fig. 3A, the reactivations often consisted of synchronous activity in many cue-responsive neurons (Fig. 3A, Supplementary Fig. 6a-b). In line with previous work, the reactivations were associated with periods of low arousal, when the pupil was constricted (Fig. 3B). This was in contrast to running and stationary periods where the animal was alert (activity during these times was defined as “other” based on baseline recording taken before training in the same session). The reactivation events persisted at similar incidence

**Figure 3:**
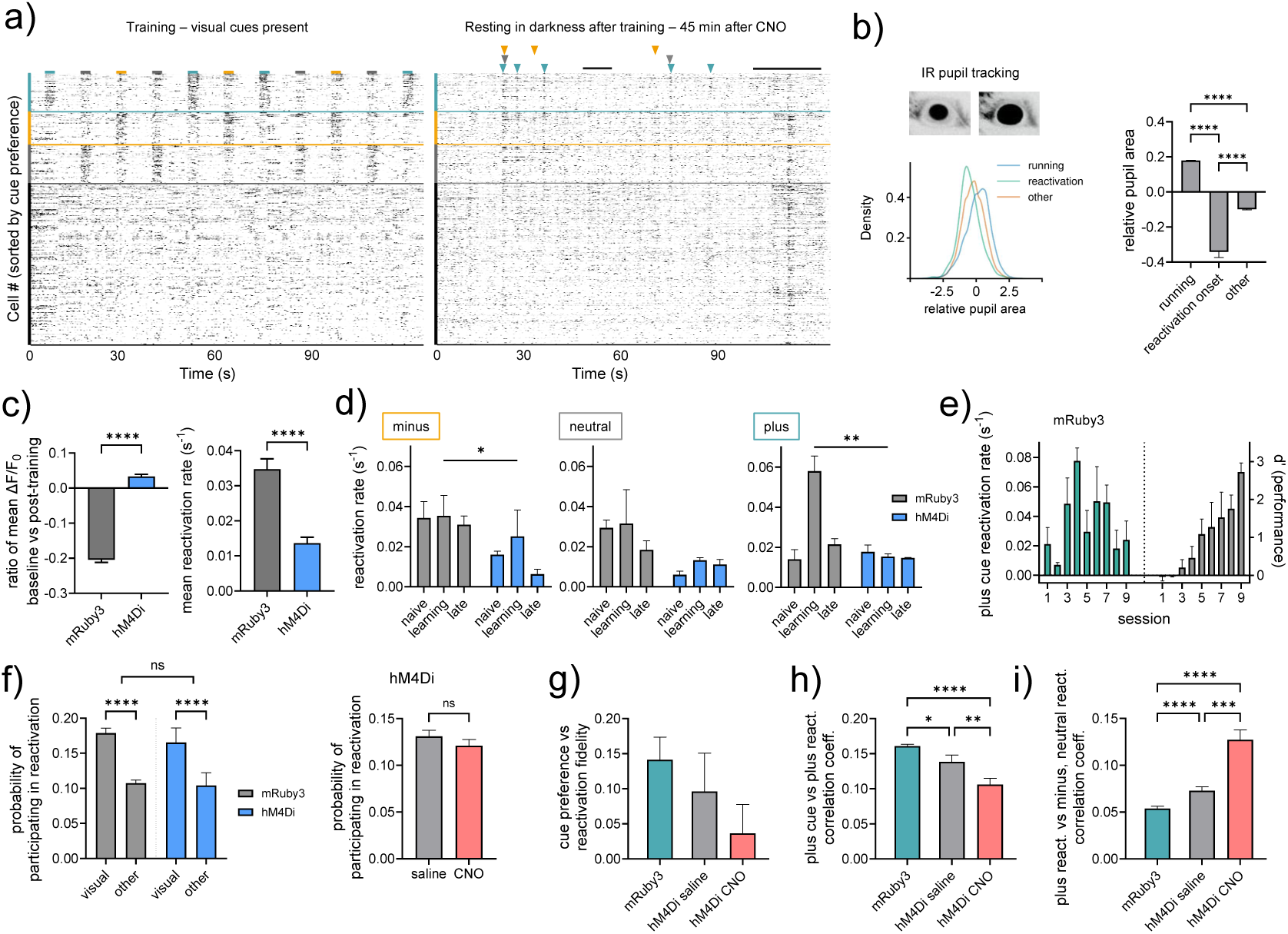
Cue-specific reactivations are attenuated by offline hM4Di activation. **a)** Raster plots from a representative session, during training (left) and post-training rest (right). Cells are sorted by cue preference. Cue identity and duration are indicated by color bars on top (left) and the corresponding identified reactivation events by color-coded arrows (right). Dark lines indicate periods of locomotion that were excluded from analysis. **b)** Pupil area was tracked by IR imaging during baseline and post-training rest recordings. Left: example images and the distribution in an example mouse of the relative pupil area during periods of running, reactivation onset, and “other” (i.e., events similar to those in baseline recordings prior to daily training), compared to the running mean across the recording. Right: quantification of the relative pupil area. One-way ANOVA, F=2524, p<0.0001 for all pairwise comparisons (Tukey’s multiple comparison test). **c)** Ratio between activity in pre-training baseline vs post-training rest (left, calculated by (rest-baseline/baseline) and mean reactivation rate per second across all cues, sessions, and animals. For both comparisons, p<0.0001, Mann-Whitney U-test, **d)** Minus, neutral, and plus cue reactivations across stages of learning. We observed a small but significant group effect for minus (p=0.04) and significant effects both of time and between groups for plus (p=0.0094, p=0.0052, generalized linear mixed-effects model). **e)** Plus cue reactivation rates for all training sessions in mRuby3 controls, and their behavioral performance. **f)** Probability of participation in reactivations across groups and functional cell types (visually responsive or “other.” In both groups of mice we observed significantly higher participation of visual cells compared to other cells, but no difference between groups. Two-way ANOVA, F (9, 67) p<0.0001 within groups, p=0.24 between groups. Within hM4Di mice, we found no significant difference between saline and CNO sessions (p=0.25, Mann-Whitney U-test). **g)** Fidelity of the cue preference versus reactivation types, defined as the ratio of reactivation participation in events of the preferred cue versus the two other cues as (preferred-other/other). **h)** Pearson correlation of plus cue presentation in training and plus reactivation events. p=0.01 (mRuby3 vs hM4Di saline), p<0.0001 (mRuby3, hM4Di CNO), p=0.009 (hM4Di saline vs CNO), Kruskall-Wallis test with Dunn’s multiple comparisons. **i)** Pearson correlation of neuronal activity during plus reactivation events and the mean of minus and neutral reactivation events. p<0.0001 (mRuby3, hM4Di saline), p<0.0001 (mRuby3, hM4Di CNO), p=0.0003 (hM4Di saline vs CNO), Kruskall-Wallis test with Dunn’s multiple comparisons. rates throughout our recording period (up to 2 hours after training) (Supplementary Fig. 6e).

To investigate how offline activity and reactivations were affected by PV+-targeted hM4Di activation, we first compared the activity of all cells in the pre-training baseline recordings with post-training rest. Only data from periods where we analyzed reactivation activity was included, i.e., not when the animal was moving or was alert. The fraction of time spent in quiet waking (i.e., not moving) was not significantly different between the groups (Supplementary Fig. 6c). We hypothesized that the inhibition of PV+ neurons would shift the population activity towards more excitation. Indeed, while mRuby3-injected mice showed reduced activity after training compared to pre-training baseline recordings, the overall activity in hM4Di-injected animals was slightly increased (Fig. 3C, left). This finding remained consistent even when we performed separate analyses of cue-responsive and other cells during training (Supplementary Fig. 7a). However, when comparing the incidences of reactivations within the same time periods, mRuby3-injected controls showed significantly higher reactivation rates than hM4Di-injected mice (Fig. 3C, right). This indicates that specifically reducing PV+ activity diminished reactivations without reducing the overall activity level.

To further explore the relationship between reactivations and consolidation, we compared the incidence rates of reactivations for each cue over time. If reactivations are indeed important for consolidation, we would expect differences in cue saliency to be mirrored in cue reactivations. Additionally, the incidence rates of these reactivations should reflect the animal’s performance [Sugden et al., 2020]. Again, we focused on the three phases of training, from the naive to the late stages. Reactivation rates were significantly higher in mRuby3-injected controls compared to hM4Di-injected mice across both salient cues (minus and plus)(Fig. 3D and E). Reactivation rates in hM4Di-injected mice did not show any significant changes across training (Fig. 3D). Notably, plus-cue reactivations were more frequent during the learning phase for mRuby3-injected controls compared to the naive and late phases. This was also evident when examining the entire training period, particularly when comparing the relative reactivation rates normalized to the non-salient neutral cue (Supplementary Fig. 7). This is in alignment with previous work [Sugden et al., 2020].

The increased reactivation rate for the “plus” cue may stem from the larger population responding to this cue as training progressed (as shown in Fig. 2C). To account for this, we first estimated the proportion of reactivation events that each cell participated in, denoted as the probability of participation in reactivations for each cell in the recorded population. Participation in reactivations was defined by cells whose activity was higher (*>* 2 s.d.) in reactivation events compared to their mean activity. We found no difference between cues or training stage (Supplementary Fig. 7c). Notably, cue-responsive cells showed higher participation compared to others in both groups of mice (Fig. 3F, left panel), but we did not detect any differences between groups or when we compared CNO and saline sessions in hM4Di mice (Fig. 3F, right panel, (Supplementary Fig. 7d)). Next, we determined the number of cells that had the greatest influence on the reactivation classification (defined as having a feature importance *>* 2 s.d. above the mean). Again, we observed no significant differences between cues, training stage, or experimental groups (Supplementary Fig. 7d).

In hM4Di-injected mice, we observed no differences in the incidence rate or cue bias of reactivations between saline and CNO sessions. Given the significant difference in behavior we had observed between these session types (Fig. 1), we performed a more detailed analysis of the content of reactivations. Our analysis focused on the plus cue responses and plus reactivations, as these accounted for the most significant learning-induced changes in mRuby3 controls. We first employed a coarse measure of specificity by calculating the fidelity between cue bias and reactivation participation across the different types. Briefly, for each session, we used the population of neurons that were tuned to the plus cue and measured the ratio of participation in plus reactivations versus minus and neutral reactivations, calculated as (plus-mean of minus and neutral)/(mean of minus and neutral). From this, we observed a strong tendency wherein mRuby3 sessions showed the highest fidelity, saline sessions in hM4Di mice showed intermediate fidelity, and CNO sessions in hM4Di mice showed very low fidelity between cue and reactivations (Fig. 3G).

This suggests that the specificity of reactivations might be compromised, as events in hM4Di mice appeared more generalized despite having the same ratio of participation from visual and other cells within the population. To investigate this further, we compared the similarity of reactivations to stimulus responses by calculating population vectors for responsive cells during the first 50 trials for each cue. These population vectors were then correlated with the population vectors of each reactivation type. Our analysis showed that the correlation between the stimulus-response and reactivations was strongest in the mRuby3 controls (Fig. 3H). While reactivation events in saline sessions from hM4Di mice showed weaker correlations than mRuby3 mice, the correlations were significantly lower during CNO sessions. Finally, when comparing reactivations of the plus cue to other reactivation events (minus, neutral, other), we found that the correlation (i.e., the similarity in reactivation patterns) was significantly stronger during CNO sessions compared to saline (Fig. 3I). The mRuby3 controls showed the lowest correlation, indicating that these reactivation events were more specific to the visual cue. Together, these results suggest that both the incidence and specificity of reactivation events were attenuated by local perturbation of the inhibitory network.

## 3 Discussion

In this work, we demonstrate that cortical reactivations in the lateral visual cortex are crucial for learning visual associations. Specifically, we show that reducing activity in the local inhibitory network during post-training rest impairs learning and significantly decreases both the incidence and specificity of reactivations while slightly increasing general spontaneous activity. During training, the cortical network remained engaged, displaying robust visual tuning responses with comparable population activity to controls. Behavioral performance improved after sessions without perturbation and within daily training runs. Overall, the most notable effect on the neural activity from reduced activity of PV+ cells was the reduction and generalization of reactivations. In contrast, control mice showed changes in response properties towards the rewarded cue, which paralleled their improved behavioral performance. These changes were reflected in reactivations in these animals, which were biased towards the rewarded cue during the training period.

Reactivations have been described for many brain regions and behavioral tasks. We have recently shown that cue-specific reactivations occur in the lateral visual cortex and affect the functional connectivity within populations of cells that respond to salient cues [Sugden et al., 2020]. These events persist for hours after training, as we also demonstrate in the current study (Supplementary Fig. 5D). To manipulate reactivation content over such extended periods without directly interfering with excitatory neurons that were active during training, we opted to perturb the activity of PV+ neurons using the DREADD system. This approach enabled two key advantages: first, the activity of the targeted neurons is reduced rather than completely silenced, and second, the perturbation can be sustained over extended periods, lasting up to 9 hours [Alexander et al., 2009, Jendryka et al., 2019]. The extensive local connectivity of PV+ neurons [Packer and Yuste, 2011] and the relatively low density of these cells in visPOR [Bjerke et al., 2021] further enhanced the likelihood that the effect of their perturbation remained relatively localized.

Using this approach, we demonstrate that learning was completely prevented, even though the manipulations occurred only *after* the training session (Fig. 1J). Previous experiments have shown that the visPOR is important for the integrity of visual association memories [Ramesh et al., 2018] and nearby cortical areas have been shown to be important for similar associative memories (e.g. Sacco and Sacchetti [2010], Grosso et al. [2015]). However, most previous work has employed relatively coarse methods of perturbing activity using cannulae-guided injections of the GABA agonist muscimol, TTX, or lesions. Our results clearly demonstrate that the learning of such associations specifically depends on an intact neural network in the visPOR in both hemispheres.

Earlier work has indicated a role for PV+ neurons in memory consolidation through effects on SWRs and their coupling to cortical spindles in the anterior cingulate cortex [Ognjanovski et al., 2017, Xia et al., 2017]. These studies show that reducing the activity of PV+ neurons in the CA1 region of the hippocampus leads to a loss of SWRs [Ognjanovski et al., 2017] and a modest reduction in coupling between cortex and hippocampus when inhibiting the activity of PV+ neurons in either area [Xia et al., 2017]. These results contrast with our findings, as we did not observe any differences in synchrony between the visPOR and hippocampus. However, unlike the prefrontal-hippocampal system, connectivity between the visPOR and CA1 is relatively sparse. Instead, the primary connectivity with the hippocampus is mediated through the entorhinal cortex [Wang et al., 2012, Estela-Pro and Burwell, 2022]. This also aligns with the somewhat weak coupling observed in our recordings (Fig. 1J). Furthermore, recent studies have shown that long-range synchrony of ripples and spindles relies on inhibitory neurons expressing somatostatin rather than PV [Abbas et al., 2018]. Additionally, the coupling between cortical activity and hippocampal ripples varies among different cell populations [Jeong et al., 2023].

Our neural activity data and behavior from hM4Di mice suggest that the cortical network remains in a naive-like state similar to that observed on the first day of training. The day-to-day fluctuations in performance indicate that the mice were well-equipped to learn, as their performance improved following training sessions with saline injections, and indeed within training sessions (Fig. 1. The cortical responses during training show that a similar proportion of the population responded to visual cues throughout the experimental period, and the distribution of cue bias matched that of the naive mRuby3 controls (Fig. 2J). Finally, when we examined the reactivation content - specifically the similarity to cue responses and among different reactivation types - we found that the sessions following saline injections exhibited an intermediate level of generalization, positioned between the pooled sessions from mRuby3 controls and CNO sessions in hM4Di injected mice (Fig. 3G-J).

A long-standing hypothesis of memory consolidation has been that the coupling of oscillations between the hippocampus and cortex initiates reactivation in relevant populations across the brain. However, despite this coupling, local regulation of activity remains crucial for controlling the fine-scale and complex firing patterns within ensembles of neurons. Moreover, recent work has suggested that increased activity of relevant populations in the cortex precedes hippocampal SWRs [Jeong et al., 2023]. When reducing the activity of PV+ neurons, we found a significant increase in the overall spontaneous activity. While not surprising in itself, this finding suggests that the overall noise within the population may prevent the emergence of fine-scale firing patterns of reactivations. It is important to note that reactivations still occurred and that the participating cells were largely consistent -primarily comprising neurons whose activity was driven by the visual stimuli.

Our data opens for a wide range of experiments to better understand memory processing. For one, while disruption of the relationship between SWRs and cortical spindles has been shown to disturb consolidation (e.g., Maingret et al. [2016]), our data suggests that this coupling may remain intact while consolidation is still disrupted. Moreover, previous data [Sugden et al., 2020] suggested that activity in the nucleus accumbens is highly correlated with reactivations in the cortex. Earlier work has indicated that learning of operant behavioral tasks that require many repetitions to achieve high accuracy, such as the one employed here, relies on consolidation and not reconsolidation in the nucleus accumbens [Hernandez et al., 2002]. How these two processes differ with respect to reactivations is unexplored. Finally, the content of reactivations and the contributions of different cell populations and types are far from understood. Our results lay the foundation for more detailed investigations by showing selective attenuation of reactivations by reducing the activity of cortical PV+ neurons.

In summary, we show that transiently reducing the activity of PV+ neurons in the offline state prevents cue-specific reactivations and learning while leaving the cortex well-equipped to learn the next day. This indicates that local inhibitory neurons are required for reactivations, which in turn are essential for memory consolidation.

## 4 Methods

### 4.1 Data reporting and statistics

No statistical methods were used to predetermine the sample size. The experimenters were not blinded during data collection. Animals from the same litters were randomly assigned to a group before surgery, and whether mice received CNO or saline on the first training day was randomized. All data was tested for normality, and appropriate statistical tests were applied. For comparisons with more than one parameter, e.g., group across time across different cues, we mainly employed a generalized linear mixed-effects model (GLMM), which accounts for shared variance and repeated measurements from the same population. If all data points were available and matched, we used RM ANOVA. All graphs in main figures represent mean±s.e.m. Analysis was performed using a combination of custom and open-access scripts in Python and Matlab. Statistical analysis and data presentation were performed using GraphPad Prism 10.

### 4.2 Experimental animals

The experiments were conducted at two locations in parallel, with similar procedures and housing conditions. The work with experimental animals that included imaging was performed at the animal facility at the Department of Bioscience, Oslo, Norway, in agreement with the guidelines for work with laboratory animals as described by the EU directive 2010/63/EU and the Norwegian Animal Welfare Act of 2010. The experiments were approved by the National Animal Research Authority of Norway (Mattilsynet, FOTS ID 14680 and 29491). The experiments with experimental animals that were used for behavioral testing and pilot experiments for imaging were performed at Beth Israel Deaconess Medical Center, Boston, USA, All animal care and experimental procedures were approved by the Beth Israel Deaconess Medical Center Institutional Animal Care and Use Committee. The animals were housed in groups of 3-6 until surgery in GM500 IVC cages, with ad libitum access to food and water. For enrichment, the cages had a running wheel, nesting material, and chew blocks made from wood.

PV-Cre mice (strain # 017320, RRID: IMSR_JAX: 017320) were originally purchased from The Jackson Laboratory and locally bred to maintain a colony. Mice of both genders were used for the behavioral experiments, but only males were used for imaging as the variability in size between individuals was smaller. This allowed us to use skull coordinates for a reliable location of virus injections and window implants, thereby reducing the overall number of animals used. At 12-15 weeks old, the mice were split into single housing and prepared for surgery. The housing room had a 12:12 hour light:dark cycle, with light on at noon and off at midnight. The ambient temperature in the housing room was kept at 21±2 °C with 25% humidity. Experiments were performed in both the light and dark phases, either in the latter half of the light phase or the first half of the dark phase.

### 4.3 Surgical procedures

The surgical procedures were similar to the ones described in [Goldey et al., 2014]. The mice were anesthetized with isoflurane (3% induction, 1-1.5% maintenance, Somnosuite, Kent Scientific) and placed on a heating pad in a stereotaxic frame (David Kopf). The depth of anesthesia was monitored throughout, and sterile Ringer’s solution heated to 37° C injected at 1-hour intervals (0.35 mL/hour). Dexamethasone (5 mg/kg, MSD Animal Health) was delivered through an intramuscular injection in the gastrocnemius, buprenorphine was administered subcutaneously (0.05 mg/kg, Individor Ltd), and local anesthetic bupivacaine (Aspen) injected in the scalp before starting further procedures. The eyes were covered with white Vaseline to prevent drying and protect them from light, and fur was shaved from behind the eyes to the upper neck region. The skin was cleaned with 70% ethanol and Jodopax Vet., before a small piece of skin covering the skull was cut away. The periosteum was removed using fine forceps and cotton swabs, and the surface was lightly scored with a scalpel. A small craniotomy was performed over the lateral visual/postrhinal cortex area of the right hemisphere using a Perfecta 300 hand-held drill (W&H) with a 0.8 mm drill-bit. Glass capillaries (OD 1.14 mm and ID 0.53 mm) pulled and beveled at a 35-40° angle were back-filled with mineral oil, and a virus injection was made using a Nanoject 3 (Drummond Scientific) of either AAV-hSyn1-Flex-mRuby3 (php. eB made in-house as described in [Grødem et al., 2023], GC 1.54 x 1013) or pAAV5-hSyn1-DIO-hM4D(Gi)-mCherry (#44362, Addgene). A total of 150 nL was injected in 30 x 5 nL steps, and the pipette was left in place for a few minutes before retraction. The exposed area was covered with KwikCast (World Precision Instruments). Next, a custom titanium head-post was attached to the skull using cyanoacrylate (Vetbond, 3M) and secured with dental acrylic C&B SuperBond (Parkell). A 3.0 mm craniotomy was then made over the left lateral visual/postrhinal cortex using a hand-held drill. The exposed cortex was flushed repeatedly with sterile Ringer’s solution kept at 37° C, and a virus injection was performed. Here, a total of 300 nL containing a 1:1 mix of AAV-hSyn-RiboL1-jGCaMP8s (php. eB made in-house ([Grødem et al., 2023]), GC 2x 1013) with either AAV-pCAG-Flex-mRuby3 or pAAV5-hSyn-DIO-hM4D(Gi)-mCherry. Custom cranial windows were prepared by attaching a 3.0 mm diameter round glass (Tower Optical) with 0.45 mm thickness to a 5.0 mm diameter glass (Warner Instruments) with 100 *µ*m thickness using Norland Optical adhesive (Thorlabs GmbH) under UV light. The cranial window was implanted and secured with C&B Superbond, and a 3D-printed light shield was attached to the head post with black dental acrylic. At the end of surgery, the mice were injected subcutaneously with meloxicam (5 mg/kg, Prion Pharma) and placed in their home cage on a heated table until fully recovered. Meloxicam injections were repeated for three days.

The coordinates used for injections relative to Bregma were −4.65 mm AP, 4.35 mm ML, and 0.4 mm ML (relative to the surface of the brain), with slight adjustments made to match the position relative to brain surface vasculature between individuals. One to two weeks after surgery, the windows were inspected under a microscope before starting behavioral training to confirm virus expression and any possible bone growth. In the case of the latter, the window was removed under isoflurane anesthesia, any bone growth or other debris cleared away, and a new window was implanted as described above. Similarly, if the expression of RiboL1-jGCaMP8s was not visible within three weeks, the cranial window was removed, and 150 nL AAV-hSyn-RiboL1-jGCaMP8s was injected.

### 4.4 Behavioral training

Two to three weeks after surgery, mice were gradually food-restricted to approximately 85% of their ad libitum body weight across four to six days. During this period, they were introduced to the behavioral apparatus on the imaging rigs across several sessions. Habituation consisted of being head-fixed on a custom 3D-printed plastic running wheel using optical posts that were mounted to the optical table, holding clamps (Standa), and modified ball joints (Thorlabs GmbH), allowing for adjustments in AP elevation. Rewards of the high-calorie milkshake Ensure were delivered by hand every 4-5 minutes. As the mice became more comfortable, the period was extended to last from 15-20 minutes up to one hour. In the final two to three sessions before introducing the operant task, the mice were shown Pavlovian visual stimuli, where the “plus” cue was shown for 3 seconds and always followed by the delivery of a small drop of Ensure. This Pavlovian training was necessary to make sure that the animals participated in the operant task from the very first session and was conducted until the mice responded to at least 50% of cue presentations.

We then began daily recording sessions. The mice were trained to discriminate full field, drifting sinusoidal gratings with a spatial frequency of 0.04 cycles per degree and a temporal frequency of 2 Hz. Each of the outcomes (delivery of 8 *µ*L of Ensure, no outcome, or 5 *µ*L of 0.1*M* quinine) was randomly assigned per animal to the different angled gratings (0°, 135° or 270°). The visual stimuli were generated using the open-source Python software PsychoPy, and synchronized with the imaging data through a parallel port and PCIe 6321 data acquisition board (National Instruments). The cues were shown on a 20” 3:4 LCD monitor (Dell, 60Hz refresh rate) or a 16” 16:9 LED monitor (Asus, 60 Hz refresh rate) positioned 25 cm from the mouse. The cues were presented for 3 seconds before a 2-second response window where licks were detected using a 3D-printed lick spout (printed in conductive filament) coupled to an MPR121 capacitive sensor (Adafruit). Licking in the response window resulted in a single delivery of Ensure or quinine at the detection of the first lick while licking outside the response window (e.g., during cue presentation) gave no outcome. Single trials were interleaved with a grey screen lasting for 5 seconds. The behavioral task was run using an Adafruit Trinket Pro and a custom circuit board run by Arduino software (similar to Sugden et al., 2020), and the delivery of Ensure and quinine was controlled by solenoid valves (#NPV-2, Clippard). Each 30-minute training session contained 54 presentations of each cue or 162 trials in total, and one session consisted of three such 30-minute runs. In the first four days of training, 5% of random “plus” trials were Pavlovian to keep the mice engaged in the task. The outcome associated with each of the cues was randomized between animals. At the end of each training session, the mice received an intraperitoneal injection of 125 *µ*L 0.9% sterile saline solution, or the same volume containing 5 mg/kg Clozapine-N-oxide (CNO) dissolved in saline (HelloBio). All animals were injected with saline and CNO on alternating days. Mice used for behavior only were placed back in their home cage and returned to the colony until the next day. Mice used in imaging or electrophysiology experiments were injected while head-fixed on the running wheel. Five minutes after the injection, the lick spout was re-positioned in front of the mice and *ad libitum* Ensure was provided for 35-40 minutes before recordings of spontaneous neural activity commenced. The behavioral performance was estimated using the discriminability index (d’) [Ramesh et al., 2018, Sugden et al., 2020], calculated as the ratio between the standardized hit rate (trials with licks to plus cue) and false alarm rate (trials with licks to minus cue).

### 4.5 Wide-field imaging

Wide-field imaging was used to monitor the expression of RiboL1-jGCaMP8s and mRuby3/hM4Di-mCherry, the quality of the cranial windows, and used daily to center the field of view to the same approximate position as previous recording days. Single images were acquired by a Canon EOS 4000D camera through a 5x Mitutoyo long working distance objective (0.14 NA) or a 16x Nikon objective (NA 0.8) in an Olympus BX-2 microscope. The light source was a xenon arc Lambda XL lamp (Sutter Instruments) with 480/545 nm and 560/635 nm filters (#39002 and #39010, Chroma).

### 4.6 Two-photon imaging

Imaging was performed at an acquisition rate of 30.9 Hz using a resonant-galvo Movable Objective Microscope (Sutter Instruments) with a MaiTai DeepSee laser (SpectraPhysics) set to a wavelength of 920 nm, or 1020 nm for reference images of hM4DI-mCherry and mRuby3. Data was collected through a Nikon 16x (NA 0.8) objective (field of view of approximately 1050x890 µm) or an Olympus 20x (NA 1.0) objective (field of view of approximately 870x700 µm). The laser output was controlled by a Pockel’s cell (302 rm, Conoptics) and set to 30-55 mW, and fluorescence was detected through bandpass filters (HQ535-50-2p and HQ610-75-2p, Chroma) by PMTs (H10770PA-40, Hamamatsu). The microscope was tilted at an angle of 4-12°, in addition to a 2-5° forward tilt of the mouse’s head adjusted by the head-fixing apparatus.

### 4.7 Electrophysiological preparation and recording

Mice initially underwent surgery for bilateral injection of virus (as described above, using either AAV-hSyn1-Flex-mRuby3 or pAAV5-hSyn1-DIO-hM4D(Gi)-mCherry), 150 nL in each hemisphere). After injections, the skin was sutured shut, and the animals were placed back in the colony. Two weeks later, a head-post was installed, and microdrives (Axona Ltd.) carrying two tetrodes each were implanted in the dorsal hippocampus and visPOR. The tetrodes were assembled using 12 µm diameter platinum-iridium (90-10%) wire (California Fine Wire), and the polyimide enamel insulation at the very ends was removed using a lighter flame. To implant the tetrodes, a small opening was made in the dura using an ophthalmologic surgery knife, and the tetrodes were implanted at AP 1.9, ML 2.1, and DV 1.1 mm (aimed at CA1 of the hippocampus) and AP 4.65, ML 4.35, and DV 0.25 mm (aimed at visPOR), relative to Bregma. Jeweler’s screws fixed to the skull served as ground electrodes, and the microdrives were secured using dental acrylic. Post-operative care was identical to the animals used for imaging.

The mice with electrophysiological implants underwent the same behavioral training as the other animals, including injections of CNO or saline on alternating days. During the habituation period of behavioral training, the microdrives were connected to the recording system to check for noise, and gradually move the electrodes in the DV plane to 0.3-0.5 mm below dura (for visPOR) and to 1.1-1.3 mm (or whenever strong 6-8 Hz theta activity was observed during running) for the hippocampus. Due to artifacts from licking, we only recorded local field potential activity in the quiet waking period after training.

Local field potential activity was recorded at a 4800 Hz sampling rate, with a low-pass filter (500 Hz) using daqUSB (Axona Ltd). The microdrives were connected to a multichannel head stage through a preamplifier. The signals were amplified 1000-3000 times using both analog and digital amplifiers, lowpass-filtered, and stored to disk for offline analysis.

### 4.8 Histology

At the end of the experiments, the animals were deeply anesthetized by an intraperitoneal injection of Euthasol (pentobarbital sodium 100mg/kg, LeVet) and perfused intracardially with 1X PBS followed by 4% paraformaldehyde in 1X PBS. The brains were dissected out and kept at 4% paraformaldehyde overnight before being transferred to a 30% sucrose solution in 1X PBS for cryoprotection. The next day, 40µm thick sections were cut with a cryostat (Leica) and collected directly to SuperFrost Plus adhesion slides. This was necessary to maintain the orientation of the cortex and midbrain sections of each slice. The sections were dried overnight before rinsing them in 1X PBS. The sections were blocked in 2% bovine serum with 0.3% Triton X-100 in 1X PBS and incubated with primary antibodies in a blocking solution for 8-14 hours. If a secondary antibody was necessary to visualize the staining, the sections were rinsed in 1X PBS and incubated with the secondary antibody for 2 hours in 1X PBS. After incubation, the sections were rinsed in 1X PBS and ddH_2_O and mounted with a mounting medium (Ibidi). The antibodies used were Alexa 594-conjugated anti-mCherry (RRID AB_2536614, Life, 1:300 dilution), chicken anti-mCherry (RRID: AB_2722769, Abcam, 1:1000 dilution), Alexa 488-conjugated anti-GFP (RRID: AB_221477, Life, 1:1000 dilution), rabbit anti-GFP (RRID: AB_2758464, Life, 1:1000 dilution), donkey anti-chicken AlexaFluor 594 (RRID: AB_2921073, Life, 1:1000 dilution) and donkey anti-rabbit AlexaFluor 488 (RRID: AB_2535792, Life, 1:1000 dilution).

Tile scans with 15% overlap were acquired using an Andor Dragonfly spinning-disc microscope with a motorized platform through a Nikon PlanApo 10X objective (0.45 NA). Images were processed to match brightness and contrast using Fiji (ImageJ).

### 4.9 Seizure analysis

Reducing inhibitory activity in the cortex could potentially give rise to epileptiform events. To investigate if this were the case in hM4Di-mCherry injected mice after CNO, we applied the methods described by Steinmetz et al. [2017]. Briefly, the events described in Steinmetz et al. appear in the LFP as large spikes with high amplitude (>1000V) with long durations (>10 ms) and in two-photon imaging data as large Ca2+ spikes throughout the entire imaging field of view. We checked all LFP recordings from all the mice using the *findpeaks* function in Matlab and detected no such events. Overall, no event in our data had an amplitude that exceeded 200µV (example traces shown in (Supplementary Fig. 2a). For imaging, we calculated the mean fluorescence intensity across the entire field of view for each frame relative to the mean intensity across all frames (Supplementary Fig. 2b). In one single animal that had a prolonged period from surgery to recordings started, we observed a strong increase in FOV-wide spiking (Supplementary Fig. 2c). These were not of similar magnitude to earlier reports, but this animal was not trained further and was not included in the data analysis. In all other animals, we detected no epileptiform events.

### 4.10 Electrophysiology analysis

Sharp-wave ripples and cortical spindles were detected using custom Matlab scripts adapted from the “Buzcode” package from the Buzsaki lab [Lab, 2024]. We used similar parameters to those described in Xia et al. [2017]. Briefly, LFPs recorded from CA1 of the hippocampus were bandpass filtered between 130-250Hz, and the amplitude was calculated using the Hilbert transform. We included events that were detected with an amplitude between 2-5 standard deviations above the mean that lasted between 15 and 200 ms. If another event occurred within 30 ms, they were merged. All ripple events were manually inspected.

Cortical spindles were detected using similar methods. Briefly, the LFPs from POR were band-pass filtered (12-15 Hz), and the amplitude filtered using the Hilbert transform. We included events that were detected with an amplitude between 2-5 standard deviations above the mean that lasted between 200 and 2000 ms. If another event occurred within 100 ms, they were merged.

To investigate synchronous events between the hippocampus and cortex, we first calculated the joint occurrence rates between SWRs and cortical spindles. Briefly, we used the timestamps for each spindle event to locate the spindle center. Any SWR event that occurred within ±25 ms of the spindle center was counted as a joint event [Maingret et al., 2016]. To control for differences in total event rates, the data was normalized to the total number of spindle events in each recording session. We also performed cross-correlations of SWRs and spindles, using similar parameters as described earlier [Xia et al., 2017]. Cross-correlations of instantaneous amplitudes of SWRs and spindles were performed in the time window ±4 sec from spindle centers, with a sliding window of 10 ms shifts.

LFP power was calculated for each frequency band by *z* -scoring. Power spectral density (PSD) was estimated by the Welch method provided by MATLAB with a window length of 1024 and a 70% overlap.

### 4.11 Image analysis

Motion correction and automatic detection of regions of interest (ROIs) were performed using Suite2p [Pachitariu et al., 2016]. All detected cell masks were manually curated based on brightness, cell mask shape, and signal-to-noise ratio. To calculate relative fluorescence Δ*F/F*_0_, traces were first corrected for neuropil contamination using *F_c_* = *F* − 0.7 ∗ *F*_neu_ + 0.7 ∗ *F*_neu, median_, where *F* represents the raw fluorescence, *F*_neu_ is the neuropil signal defined by Suite2p as the fluorescence in a surrounding region around the ROI, and *F*_neu, median_ is the median value of that signal (adapted from [Goltstein et al., 2018]). Δ*F/F*_0_ was then calculated as (*F_c_* − *F*_0_)*/F*_0_, where *F*_0_ is the running median (with a window size of 600s) of the neuropil-corrected trace *F_c_*. The OASIS algorithm was used to deconvolve the Δ*F/F*_0_ traces [Friedrich et al., 2017].

### 4.12 Quantification of visual responses

To determine whether a cell was responsive to a stimulus, we first binned the trials into 1-second intervals starting from the stimulus onset. For each bin, we computed the relative change in the median raw neuropil-corrected fluorescence (*F_c_*) compared to a 1-second baseline period prior to stimulus onset. For each stimulus type (plus, minus, neutral), we applied a Wilcoxon signed-rank test to assess whether there was a significant increase in either the first or second bin. We set *α* = 0.01 and applied a Bonferroni correction to account for multiple comparisons across cells, stimulus types, and bins. A cell was considered responsive if it showed a significant response in one of the two bins. If a cell was responsive to more than one stimulus, we assigned it to the stimulus with the highest significant response. To selectively analyse cue-elicited responses not affected by outcome, we performed the same analysis but only included data from the first second of cue presentation, or until the first lick within this time.

### 4.13 Cell matching across sessions

We developed a custom script to create pairwise mappings between the ROI IDs of sessions captured from the same FOV. To map the cells of two sessions, we began by calculating a similarity transform between the mean images of the FOVs of the recordings, incorporating rotation, translation, and uniform scaling [Reddy and Chatterji, 1996]. This transform was then applied to the binary cell masks of the second session, followed by re-binarization of the masks using a threshold of 0.2. To identify potential cell pairs, we calculated the Dice similarity coefficients between the cell masks of the two sessions. A cell from the first session was mapped to the cell in the second session with the highest Dice score, provided that the score exceeded 0.5 and the gap to the next highest candidate was at least 0.15. These criteria allowed us to establish a mapping table between the two sessions.

Mapping of the various sessions began by selecting one session as the reference, and then all subsequent sessions were mapped to the reference. After completing this process, we selected the next session as the reference and repeated the mapping for all other sessions, excluding those previously used as references. During this iterative mapping process, many cells were already mapped indirectly (e.g., if two cells from different sessions were mapped to the same cell in the reference session, they were considered mapped to each other). We focused our efforts on mapping cells that had not yet been matched. Ultimately, we compiled a coherent mapping table that included all sessions for a given subject from the same FOV. To ensure the accuracy of our mappings, we generated cell maps for all groups and performed a manual quality check, excluding cells from the mapping if they did not exhibit sufficient similarity. The analysis of single-cell responses across training (Fig. 2D) was mainly observational and not quantified. We selected example cells from both populations that very clearly represented the overall population, where many neurons from mRuby3 controls were dynamic and shifted their tuning towards the rewarded cue, while the neurons from hM4Di injected mice were very stable over time and unaffected by daily training.

### 4.14 Reactivations

Reactivations were predicted using machine learning-based classification, using the frames as samples and the cells as variables in the matrix of independent variables. The samples were classified into one of the following classes based on the activities in the different frames: plus, minus, or neutral reactivations, or “other”, where the “other” class consisted of the samples in the baseline run (run 0). The inter-stimulus intervals in run 1, 2 and 3 were excluded from the analysis.

Initially, two different approaches for validation were compared: leave-one-run-out crossvalidation and test set validation, where the latter was found to be the most suitable approach for our study. The classification models were calibrated using a training set consisting of 70% of the samples and validated using the remaining 30%, which were kept as a separate test set. The final prediction of reactivations in the post-training rest (runs 9 and 10) was then based on the same model (trained using 70% of the samples). In order to choose the best classification method for this study, the following classification methods were compared: Multiclass perceptron [McCulloch and Pitts, 1943], logistic regression [Berkson, 1944, Nguyen et al., 2024], support vector machine [Cortes and Vapnik, 1995], Random Forest [Ho, 1998], quadratic discriminant analysis (QDA) [Hastie et al., 2009], Gaussian Naïve Bayes classification [Hand and Yu, 2001], Ada Boost [Freund and Schapire, 1997], Least absolute shrinkage and selection operator (LASSO) [Tibshirani, 1996] and Elastic net [Zou and Hastie, 2005]. In an initial comparison using a few randomly selected sessions from mRuby3 injected control animals, Random Forest, QDA, and Logistic Regression were evaluated as the top three classification methods, taking both classification accuracy and computational demand into consideration. A more detailed comparison of these three methods using all animals and sessions identified Random Forest as the most suitable classification method (Supplementary Fig. 7c). All classifications except those done using Random Forest were based on data standardized to a mean of zero and a standard deviation of one.

All classification analyses were carried out using standard modules from Scikit-learn (https://scikit-learn.org/stable/) in Python 3.11.3. The hyper-parameters of all classification methods were tuned in the training stage using exhaustive grid search (GridSearchCV from Scikit-learn) with 5-fold cross-validation.

Training and predictions were carried out per frame using the deconvolved traces. We applied a rolling maximum filter (∼ 340 ms) to those traces, similar to Nguyen et al. [2024], to account for the different timings of the individual cells. In the training phase, only every second frame was used in order to limit the computational demand, while the final predictions (for runs 9 and 10) were done for all frames. Following the prediction of reactivations using classification, the results were filtered to retain only predictions where the mice were not running. This was done in order to avoid confusing the activation of cells due to running with those due to preceding stimuli. Any period where the mouse had a running speed above 0.1 cm/sec was removed, including the 10 seconds prior and after [Sugden et al., 2020]. The results were further filtered to retain only reactivations occurring during highly synchronous events within stimulus-responsive neurons. Similar to Nguyen et al. [2024], those events were defined as frames in which the average activity of at least one responsive cell group (plus, minus, neutral) was above 3 s.d. above the mean of that group.

### 4.15 Correlations

To study the correlation between reactivation events and stimulus responses during training, we first constructed a response template for each stimulus type. Using the deconvolved trace, we applied a rolling maximum filter (∼ 340 ms) and averaged the activity of each cell over a one-second period following stimulus onset across trials to get a population vector for each stimulus type. Next, we pre-processed the activity during the spontaneous runs in the same way. The stimulus templates were then correlated with all identified reactivation events during those runs and filtered to remove running and to include only highly synchronous events (as described above). Additionally, we computed correlations with all other time points when the animal was at rest (defined as running speed below 0.1 cm/sec for at least 10 seconds), excluding reactivation events and events predicted by the classifier that did not meet the reactivation criteria, as well as five frames before and after these events.

## 5 Data and code availability

The data sets are available upon reasonable request and will be made public through the Norwegian Research Infrastructure Services. All custom code used for analysis will be made available through www.github.com.

## 6 Author contributions

K.K.L. and A.S. conceived of the project. K.K.L., A.S., M.L.A and M.F. designed the experiments., S.G. made viral constructs and assisted with co-expression experiments, K.K.L. performed surgeries, K.K.L. and I.N.N. performed the experiments, with assistance from I.S. on behavior. F.S.R. analyzed two-photon data with assistance from K.K.L. and I.N.N., and F.S.R. and K.T. designed and performed the classification of reactivations. K.K.L., T.H. and A.M.S. analyzed LFP recordings. K.K.L. wrote the first draft of the manuscript with assistance from I.N.N. and F.S.R..

## 7 Acknowledgments

We thank Sondre Jordbræk for assistance with detection of SWRs and cortical spindles, Dr. Yoav Livneh for useful discussions regarding co-expression of viruses and CNO injections, Osama Alturkistani for assistance with surgeries for pilot experiments, Dr. Alessio Buccino for assistance with the behavior apparatus, and Nghia Nguyen for useful feedback to the manuscript. This work was supported by The Research Council of Norway (grant nr 616022 to K.K.L., 248828, 250259 and 309547 to M.F. and A.M.S, 325892 to T.H.), the University of Oslo’s strategic research initiative CINPLA, and the European Union’s Horizon 2020 research and innovation programme under the Marie Skłodowska-Curie grant agreement nr 945371 to A.M.S.).

## 8 Supplementary information

The supplementary information contains supplementary figures S1-7.

**Supplementary Figure 1.**
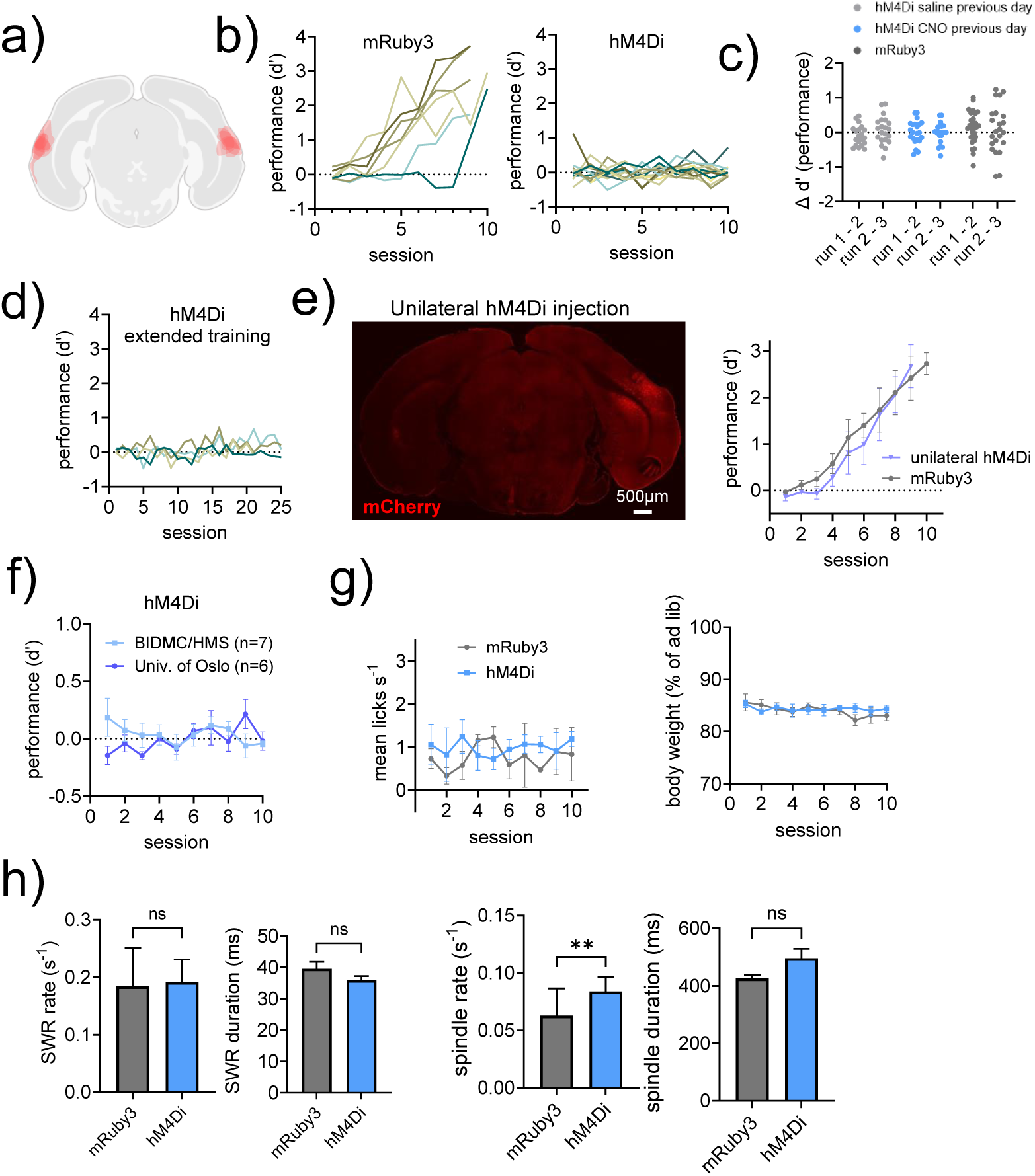
a) Spread of hM4Di-mCherry, drawn for 7 mice with bilateral injections onto a coronal section. b) Learning curves for individual animals. c) Within session change in performance for example animals. d) A subset of hM4Di-injected mice were trained for a substantial period beyond the normal experimental time frame. There was no development in the performance of these mice. e) Unilateral injection of hM4Di in visPOR did not affect learning performance. Histology example of unilateral AAV5.hSyn.DIO-hM4Di-mCherry injection and learning curve compared to mRuby3 injected mice, n=4 (unilateral) and 7 (mRuby3). f) Similar results from behavioral training was reproduced in two labs. g) We did not detect significant differences between groups for the mean number of licks per second or daily pre-training body weight. h) No significant differences in the incidence rate or duration of sharp-wave ripples were found. We detected a significant difference in the incidence rate of cortical spindles between groups (Mann-Whitney U-test, p=0.0016. n=20 (hM4Di), and 19 (mRuby3) sessions from 2 and 3 mice, respectively), but not for duration of spindles.

**Supplementary Figure 2.**
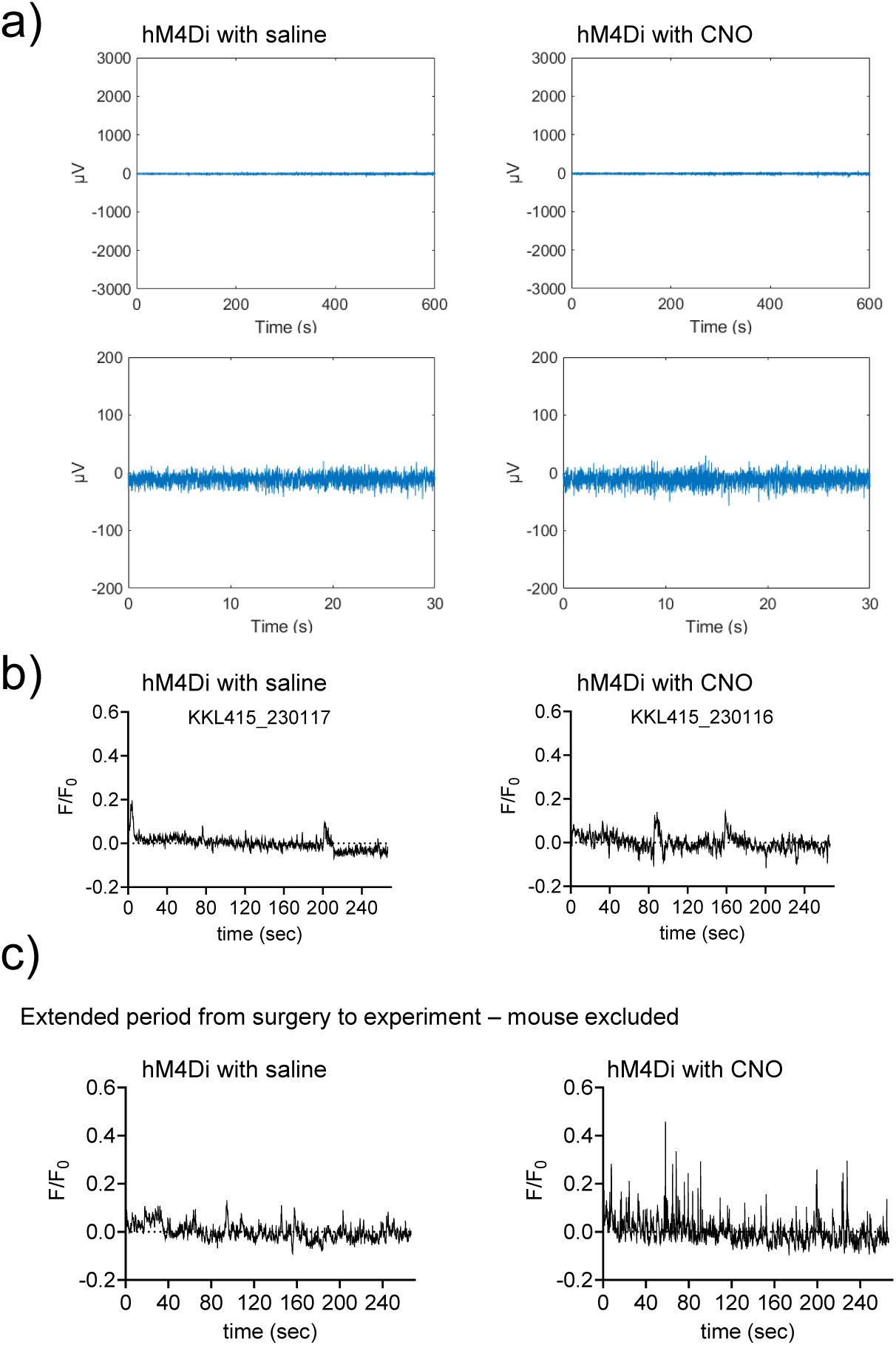
a) Representative LFP traces from visPOR in a hM4Di injected mouse during post-training rest with either saline (left) or CNO (right). We did not detect any seizure-like events in any LFP recording. Traces are shown across 10 minutes (upper panels) and 30 seconds. b) Changes in fluorescence across the field-of-view, normalized to the mean intensity. We did not detect abnormal events in any mouse used in the data presented in the main figures. c) Example from the single mouse that was excluded from analysis of reactivation events. Due to bone growth we replaced the cranial window twice before the experiment started, leading to a very prolonged period (12 weeks) from virus injections to recording. Here, we did not observe activity as described in Steinmetz et al (2017), but a high event rate of transient peaks in fluorescence.

**Supplementary Figure 3.**
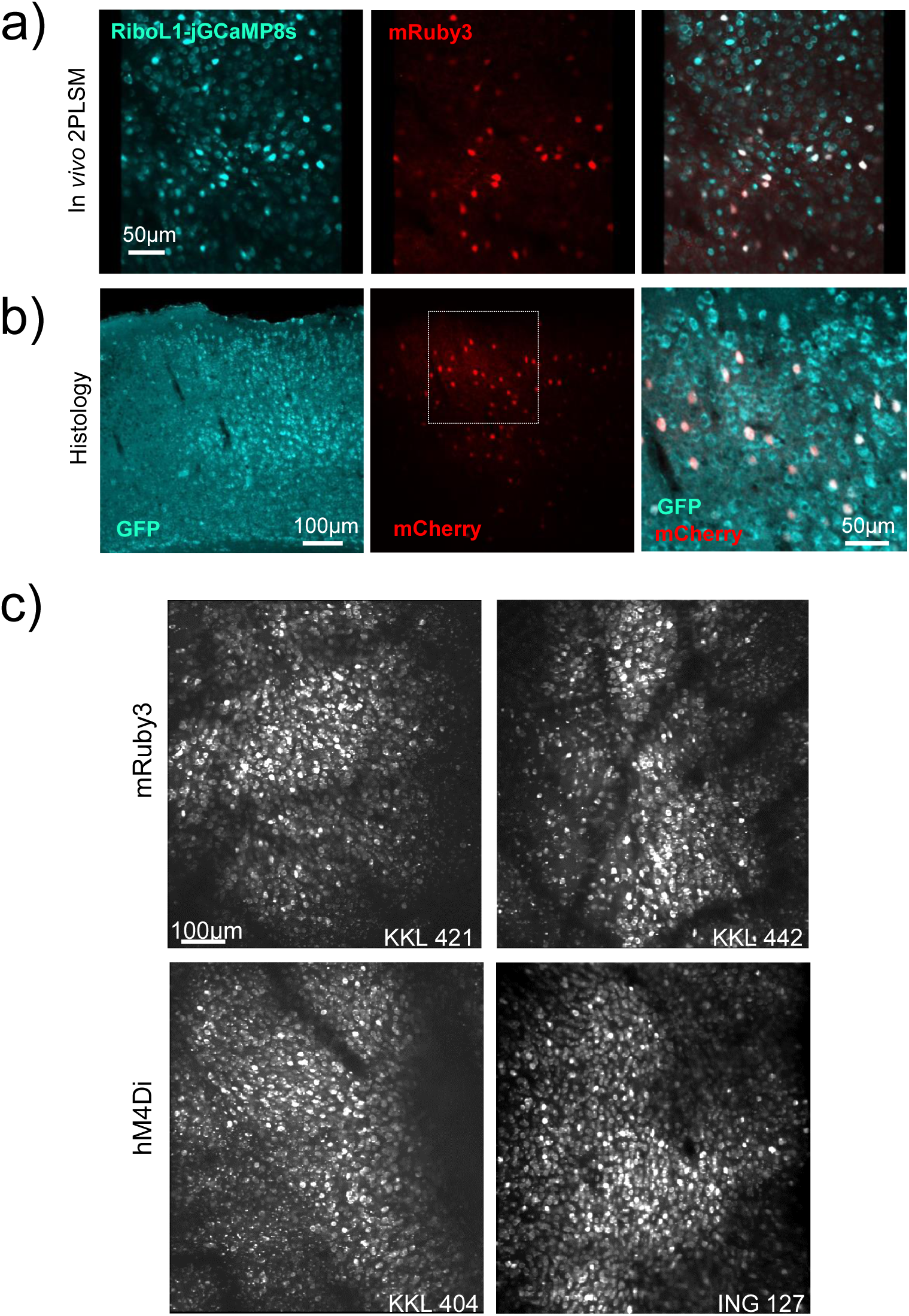
a) *In vivo* two-color 2PLSM imaging showing co-expression between mRuby3 and RiboL1-jGCaMP8s. b) Representative *post-mortem* histology showing the expression of RiboL1-jGCaMP8s (identified by GFP) and DIO-hM4Di-mCherry. c) Representative fields of view from two-photon imaging. Images exported from Suite2p as average intensity images across one imaging session.

**Supplementary Figure 4.**
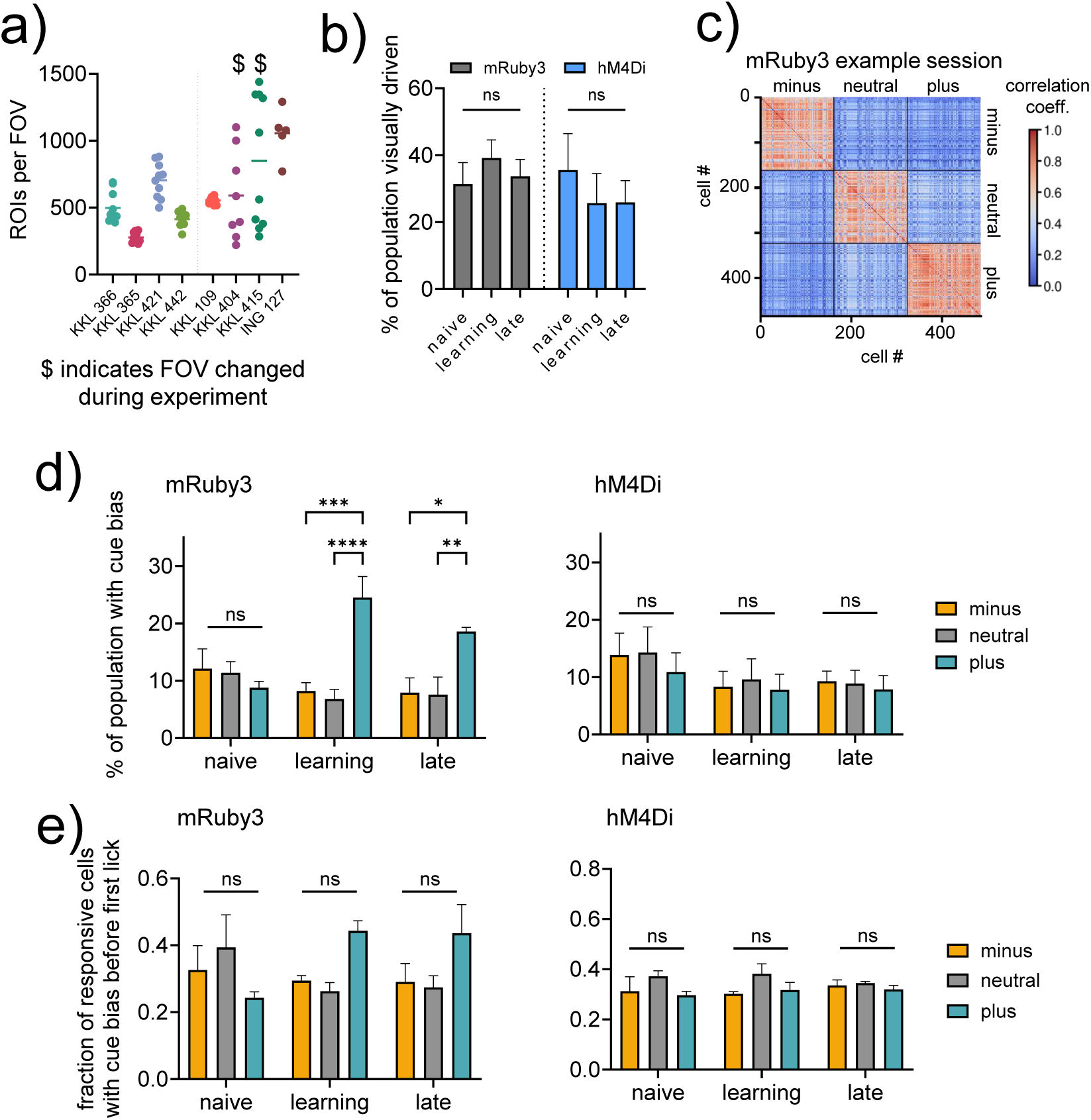
a) The number of ROIs from each imaging session per animal after manual sorting. In two mice we changed the field of view (indicated by “$”) during the experiment because of partial bone growth. b) Quantification of the mean percentage of the population responding to any visual cue. We observed no significant differences between groups. GLMM with multiple comparisons, p=0.76 (naive), p=0.26 (learning), p=0.4 (late), nor within groups across time (mRuby3 p=0.14 naive vs learning, p=0.71 naive vs late, p=0.4 learning vs late, hM4Di p=0.1 naive vs learning, p=0.28 naive vs late, p=0.99 learning vs late), c) Pairwise correlations of all cells defined as visually responsive in two consecutive example sessions, sorted by cue preference. d) Related to Fig 2C, but showing the percentage of cells with cue bias of the entire population. Two-way ANOVA with Tukey’s multiple comparisons test. F (4, 27), p=0.0002 (minus vs plus, “learning”), p<0.0001 (neutral vs plus, “learning”), p= 0.011 (minus vs plus “late”), p= 0.0088 (neutral vs plus, “late”). For hM4Di mice there were no significant differences. e) Similar to d but only including cue responses in the first sec of stimulus or before the first lick. No significant differences detected across time or group.

**Supplementary Figure 5.**
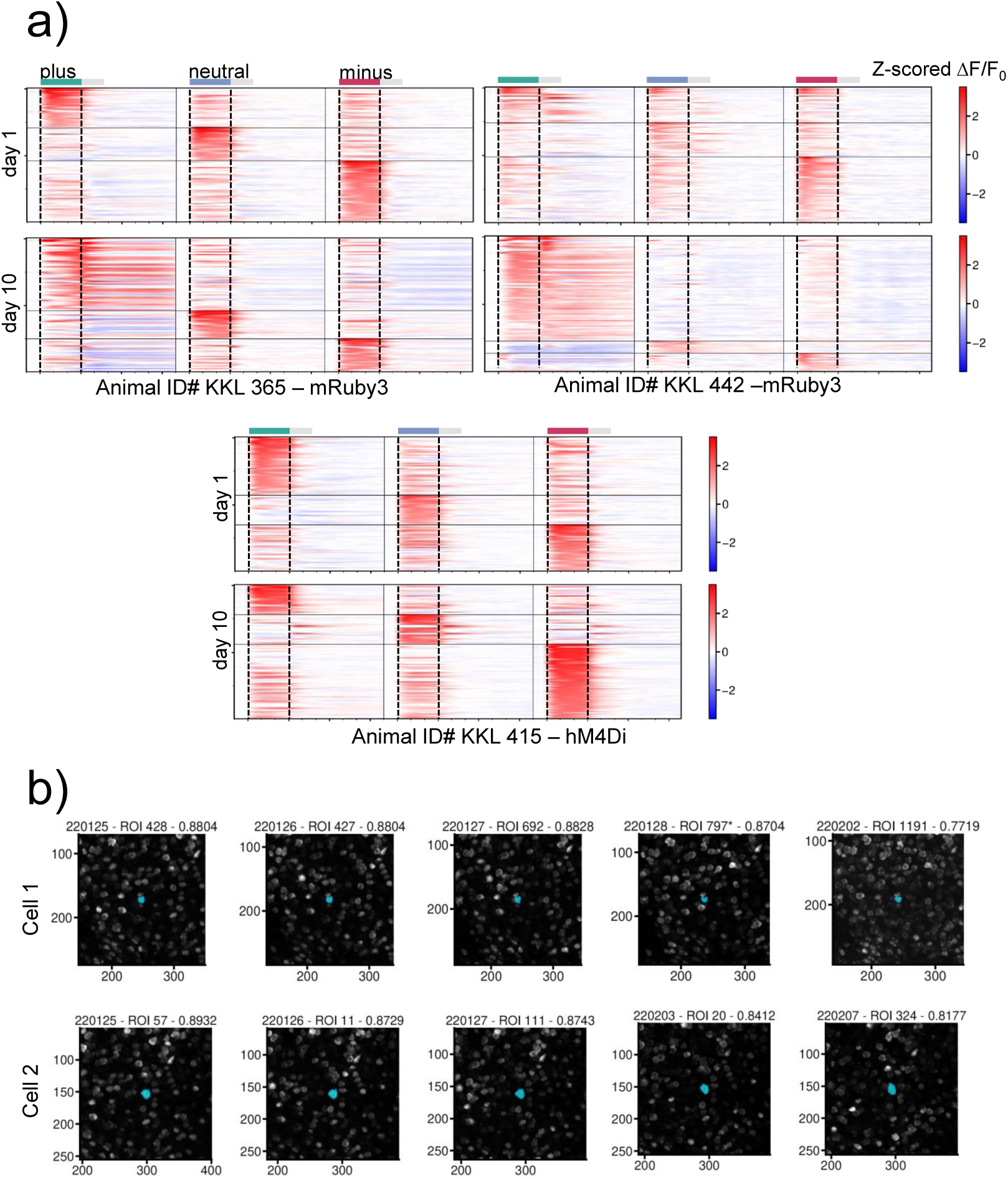
a) Examples related to Figure 2B. We observed a similar trend in all mRuby3 injected animals, where the population responded stronger to the “plus” cue with training. No such development was seen in hM4Di injected mice treated with CNO. b) Two examples of same-cell tracking across the experiment. Numbers indicate the date of the experiment, ROI number (from Suite2p) and Dice score.

**Supplementary Figure 6.**
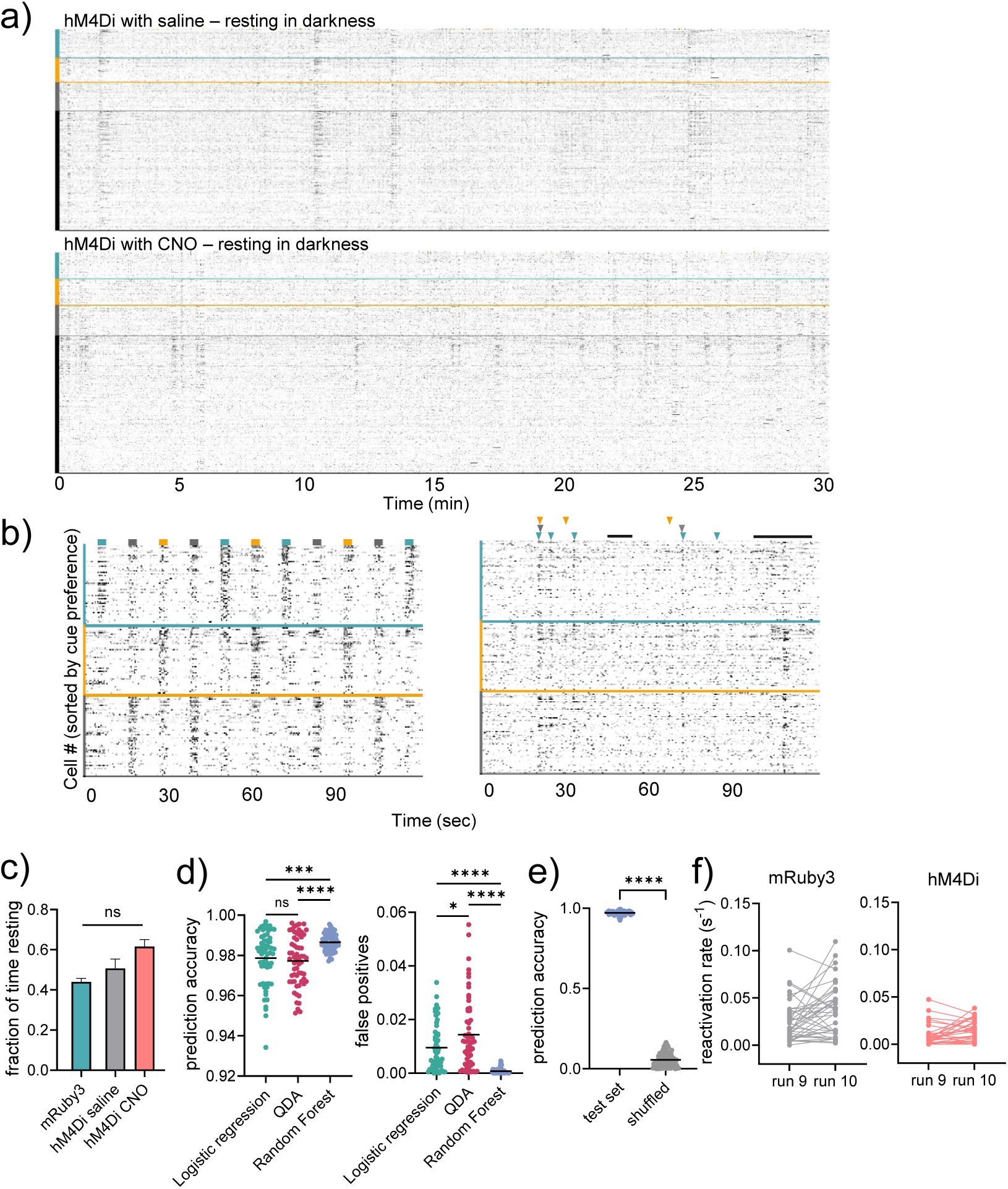
a) Representative raster plots of 30-minute recordings during post-training rest from consecutive days in the same hM4Di injected animals, one with saline and one with CNO. Cells are sorted relative to their cue preference. b) Related to Fig 3A but only showing stimulus responses and cue reactivations in cells defined as visually responsive. c) Fraction of time in post-training rest recordings spent resting. GLMM, no significant differences between groups. d) Prediction accuracy and false positive fractions using three different classification methods. Random Forest performed better when compared to Logistic regression and QDA. One-way ANOVA with Tukey’s multiple comparisons test for prediction accuracy F=12.6. p=0.75 (Log.Reg vs QDA), p=0.0003 (Log-Reg vs Random Forest), p<0.0001 (QDA vs Random Forest). For False positives, F=33.9, p=0.01 (Log.Reg. vs QDA), p<0.0001 (Log.Reg. and QDA vs Random Forest). e) prediction accuracy of visual stimuli in the test set compared to shuffled data. Unpaired t-test, p<0.0001. f) Reactivations persisted through our daily recording period (up to 2 hours after training). Example sessions from two mRuby3 mice and two hM4Di mice. “Run 9” and “Run 10” indicate the two 30-minute recordings that followed daily training, saline or CNO injections, and 45 minutes of *ad libitum* Ensure consumption, respectively.

**Supplementary Figure 7.**
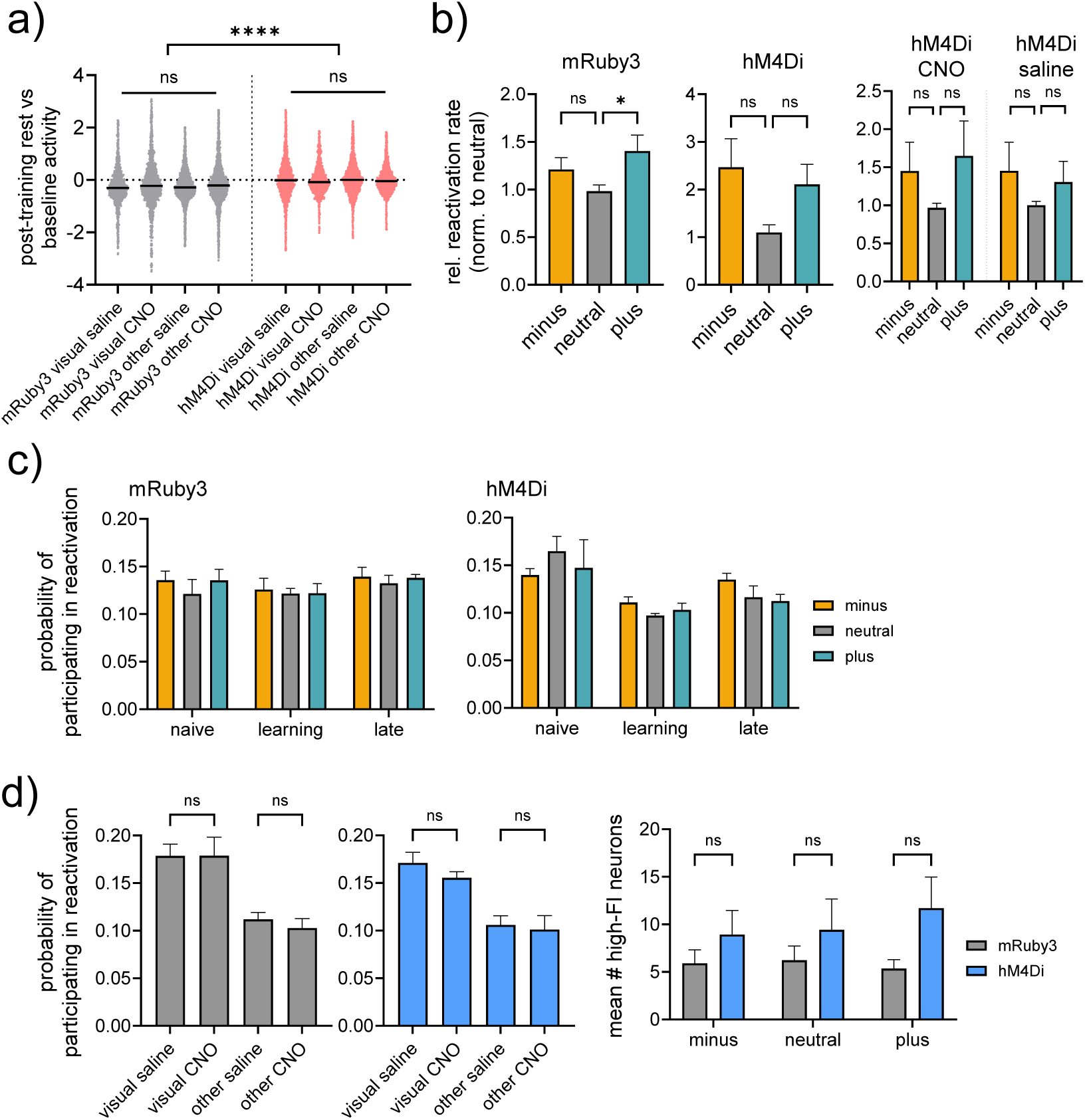
a) Relative activity in post-training rest compared to baseline, related to Figure 3C. Within mRuby3 and hM4Di groups we found no significant differences, but across groups we observed significant difference in ratios within all cell categories (p<0.0001, GLMM with multiple comparisons. b) Relative reactivation rates, normalized to the neutral reactivations in the same session. The relative rate for plus reactivations was significantly higher in mRuby3 control mice (Repeated Measures ANOVA with Dunnett’s multiple comparisons test for paired data, p=0.08 for minus vs neutral, 0.03 for plus vs neutral). For hM4Di there were no significant differences across days, or within saline or CNO sessions (right panel), c) Participation in minus-, neutral –or plus reactivations at different training stages, as a probability based on the mean participation rate per cell. d) Left: Participation in plus reactivations across visually responsive and other cells, sorted by group (mRuby3 or hM4Di) and treatment (saline or CNO). No significant differences were observed either within or between groups. Right: The mean number of cells with high feature importance for each visual cue. Two-way ANOVA, F(2,12) revealed a significant difference between groups across all conditions (p=0.01) but not for each individual cue.

